# Struvite fertiliser offers a sustainable solution for recycling of phosphorus for crop growth without altering the soil resistome

**DOI:** 10.1101/2025.10.06.680669

**Authors:** Katie Lawther, Nicholas J Dimonaco, Brett Greer, Paul N Williams, Jason P Chin, Rebecca L Hall, John W McGrath, Sharon A Huws

**Author notes:** **Correspondence: Professor Sharon Huws,** School of Biological Sciences/Institute for Global Food Security, Queen’s University Belfast, 19 Chlorine Gardens, Belfast, UK. BT9 5DL.; Dr Katie Lawther, School of Biological Sciences/Institute for Global Food Security, Queen’s University Belfast, 19 Chlorine Gardens, Belfast, UK. BT9 5DL.

## Abstract

**Background:** The reserves of rock phosphate, used to produce fertilisers, are being exhausted by the ever-increasing pressure to meet the food demands of a growing world population. Sustainable alternatives are urgently needed, and struvite, a renewable phosphorus fertiliser recovered from waste streams such as animal manure, offers a promising solution. However, other manure-derived fertilisers have been associated with the promotion of antimicrobial resistance (AMR) in soil, raising concerns about whether struvite might pose similar risks. Despite this, the impact of struvite on the soil resistome remains poorly understood.

**Methods:** This study assesses the AMR risk associated with Xiamen struvite (XIA), produced from piggery wastewater, compared with two commercial fertilisers: Crystal Green (CG) and Triple Super Phosphate (TSP), and an untreated control soil. Following a three-month fertilisation period, rhizospheric soils were analysed using mass spectrometry to investigate antimicrobial residues in the soil, and shotgun metagenomics and culture-based assays, and bacterial isolate characterisation to identify bacterial AMR linked to the treatments. Notably, this is the first study to apply culturing methods to investigate struvite’s impact on AMR.

**Results and conclusions:** While XIA struvite soil fertilisation led to an increase in total culturable bacterial load, (CFU/g), there was no evidence of antimicrobial residues detected in soil, as confirmed by mass spectrometry. Metagenomic profiling further revealed no significant alterations to antimicrobial resistance genes (ARGs) across all treatments, including CG, TSP, and untreated controls. Culture-based analysis supported these findings, with tetracycline-resistant bacteria undetected and only minimal resistance observed to erythromycin, florfenicol, and neomycin. Although resistance levels to amoxicillin, streptomycin, and trimethoprim were higher in XIA-treated soils, this appeared to reflect a broader increase in microbial load rather than selective enrichment of resistant strains. These data suggest that piggery-derived struvite does not impose a clear AMR risk in soil and performs similarly to conventional fertilisers. While it appears to increase total bacterial load, there is no evidence of selective enrichment for AMR bacteria. However, variability in antimicrobial and ARG content across studies highlights the need for standardised production specifications for struvites to ensure safety and consistency. This study provides the most comprehensive assessment to date of struvite’s AMR implications, supporting its potential as a safe, sustainable fertiliser and contributing valuable insights to inform regulatory guidance.

## Introduction

The growing human population is demanding ever more food, with crop production being reliant on nutrient input, such as nitrogen, potassium and phosphorus (Ågren *et al.,* 2012). Nonetheless, our phosphorus fertiliser reserves are depleting, thus threatening our food security. The primary readily available source of phosphorus (P) for fertiliser production is phosphate rock, and estimates indicate that domestic supplies in major producer countries, such as China (48% of global production), Morocco (11%), and the USA (10%), could be depleted within 50 years (under the assumption that no new phosphate rock reserves are uncovered, technological advances in mining are not made and production continues at its current rate). Notably, 85% of global phosphate rock reserves are concentrated in just five areas, with Morocco and Western Sahara holding the majority (70%) (Cordell, 2008; Ott and Rechberger, 2012; Jasinski, 2021). As a result, many nations rely on P imports for food production, raising the potential of food insecurity. For instance, the EU (European Union) has a phosphorus import reliance of around 84% and an end-of-life recycling input rate (EoL-RIR), this is the proportion of demand met by secondary raw materials, of just 17% (European Commission, 2020). Additionally, the quality of the remaining phosphate rock reserves is declining, with higher levels of impurities and heavy metals, resulting in increased processing costs (Cordell and White, 2011; Cordell and White, 2013; Blank, 2012; Ar, 2009). For example, cadmium, the toxic heavy metal, can be present in high levels in rock phosphate and fertilisers, negatively affecting both soil fertility and crops, and can accumulate in the food chain posing a serious threat to human health (Mar and Okazaki, 2012; Niño-Savala *et al.,* 2019).

Today the majority of the phosphorus in sewage, manure, food waste, industrial slags and other waste streams, is lost to the environment and not recycled or recovered, leading to a broken P cycle (Ohtake and Tsuneda, 2019). Mined phosphate rock (synthetic) fertilisers when applied in excess can also cause P runoff from agricultural soil, which results in phosphorus accumulation in aquatic systems, leading to eutrophication with subsequent harmful algal bloom formation and associated public health risks (Kleinman *et al.,* 2011; Oelkers and Valsami-Jones, 2008; Reid et al., 2024). Given both supply chain and rock quality concerns, alongside an ever-expanding global population and the associated pressures on food production, the development of sustainably derived fertilisers including those produced via P recovery from waste, has sought to offset agricultural reliance on mined P rock.

One potential candidate for a renewable P fertiliser is waste recovered struvite (magnesium ammonium phosphate: NH4MgPO4·6H2O). Struvite is formed through crystallisation and can be produced from several wastewater streams including human urine, anaerobically digested dairy manure and swine wastewater (Rahman et al., 2014; Chen et al., 2017; Etter et al., 2011; Uludag-Demirer et al., 2005). Commercial struvite crystallisation has been able to recover 93% of P from wastewaters and 40% of nitrogen, therefore, it could potentially meet part of the increasing phosphorus demands and supplement the finite natural P resources (Rahman *et al.,* 2011; Degryse *et al.,* 2017). Advantages of recovering P fertilisers commercially include reducing biosolids in waste and a reduction in chemical sludge, leading to decreased treatment plant maintenance, lowering the P content of waste and the sale of the fertilisers produced can offset operational costs (Ostara, 2021). Struvite can be successfully used as a highly efficient fertiliser for tree seedlings, ornamentals, flower boards, garden and turf grass and vegetables (Rahman *et al.,* 2014). Struvite fertilisers have been shown to perform equally to the widely used chemical phosphate fertiliser Triple Super Phosphate (TSP) in terms of plant growth and P-bioavailability (Hall *et al.,* 2020). Struvite fertilisers have also been shown to have lower concentrations of toxic metals than TSP (Hall *et al.,* 2020). Due to slow-release properties of struvite, low amounts of P are lost in runoff (1.9% of P dose), including during rainfall, reducing the risk of eutrophication in surface water (Everaert *et al.,* 2018). An example of commercial struvite fertiliser production methods is Ostara’s Pearl® process used to produce Crystal Green, briefly this includes the addition of magnesium to wastewater in a pH-controlled method, the struvite then precipitates out from the water before being heated and dried (Evoqua Water Technologies, 2021). Due to this wastewater origin, commercial struvite production is required to operate under strict requirements including low potentially toxic elements i.e. heavy metals (Muys et al., 2021). Other methods of struvite production can include the addition of sodium hydroxide to increase the pH of wastewater above 8, this method was used to produce a struvite from piggery wastewater, designated Xiamen struvite (Ye *et al*., 2010).

Fertilisers produced from human or animal derived waste streams are not however without issue. Fertiliser production from such waste steams has the potential to introduce antibiotic residues that create selection pressures in soil, promoting the proliferation of resistant microbes, including pathogens (Udikovic-Kolic et al., 2014). These fertilisers may also act as sources of antimicrobial resistant (AMR) bacteria and antibiotic resistance genes (ARGs), which can colonise soil and transfer resistance to native microbes (Heuer and Smalla, 2007; Heuer et al., 2011; Wichmann et al., 2014).

Concerns have also been noted regarding the presence of ARGs and antibiotic residues contained within struvite fertilisers which may contribute to the spread of AMR (Chen et al., 2017; Woldeyohannis and Desta, 2024). Recent studies investigating urine derived struvites showed that the material contained ARGs conferring resistance to aminoglycosides, carbapenems, chloramphenicol, and erythromycin, with bacterial carriers such as *Acinetobacter*, *Aeromonas*, and *Enterococcus* spp. (Woldeyohannis and Desta, 2024). In the case of phosphorus-rich wastewater-derived struvite and livestock wastewater-derived struvite, high concentrations of antibiotics have also been reported, with end product antibiotic residue concentrations being influenced by pH, dissolved organic matter, and struvite crystallisation methods (Gao et al., 2023; Lou et al., 2018). The application of struvite in agricultural soils also raises concerns about its role in ARG proliferation. Research demonstrates that struvite amendments increase the genetic diversity of ARG cassettes in both rhizosphere and phyllosphere environments, with class 1 integrons being the most prevalent (An et al., 2018). A study investigating the effects of struvite, sourced from swine wastewater produced by an anaerobic digester, on the soil resistome post-application concluded that the struvite enhanced the soil resistome, particularly with respect to multidrug resistance (MDR) and resistance to aminoglycosides, β-lactams, macrolides, lincosamides and streptogramin B (MLSB), sulphonamides and tetracyclines (Chen *et al.,* 2017). Controversially, due to these findings, in 2019 the European Commission expressed concern about the use of struvite and stated that further research was required to investigate its impact on the soil resistome (De Boer *et al.,* 2018; European Commission, 2018). More recently in 2022, the EU updated their rules on fertilisers, allowing use of precipitated phosphate salts and derivatives (‘struvites’), including those recovered from waste streams (European Commission, 2021).

Given the potential of struvite as a fertiliser, this study investigates how struvite produced from piggery wastewater affects the soil and compares its impact to that of commercially available fertilisers, including triple superphosphate (TSP; Goulding Fertiliser NI) and Crystal Green® struvite derived from human waste streams (Ostara, 2021).

## Methods

### Pot plant experiment

The Mineral Gley soil used in this study was collected from a 3-month long pot experiment performed by Hall *et al.,* (2020). Mineral Gley soil was collected from the field boundary on unfertilised soil from a long-term grass rotation at Garvagh, Northern Ireland (NI) (Grid reference: NW022776). Humic-gley soil was used, with an organic sandy texture and a of pH 5.4 and the following characteristics: P 1.7 mg/g, Al 70 mg/g, Fe 31 mg/g, Mg 13 mg/g, and Ca 2.0 mg/g (Hall et al., 2020). Pots containing 5kg of soil (30 x 25cm) were planted with Perennial ryegrass, three replicates (pots) per condition. Full perennial ryegrass (*Lolium perenne*) germination procedures are detailed in experiment A by Hall *et al.,* (2020). Briefly, Perennial ryegrass seeds were germinated within a greenhouse in rock wool in a hydroponic growth system, the germinated seedlings were then transplanted after one month.

The experimental design included 4 treatments: unfertilised control (UNT), a commercially available struvite from municipal waste streams (Crystal Green, CG), a novel/not commercially available struvite produced from piggery wastewater (XIA) and finally Triple Super Phosphate (TSP) which is a rock P based fertiliser and currently considered the gold standard of P fertilisers (Table 1). The different fertilisers were applied to the top layer of soil and homogenous distribution was achieved through hand mixing, left to equilibrate for 4 days, and then each condition potted in triplicate. Assisted heat and lighting was supplied from 8am-8 pm to replicate temperate growing conditions, light levels and a soil water holding capacity of 65% were also maintained. Time 0 hr soil was collected immediately following the application of the different fertilisers (Table 1) and homogenous distribution (soil mixing). Following a 3-month growth period rhizospheric soil was collected from each pot plant (Time End sample). At each sampling event ∼1g of soil was collected from each pot and stored immediately at - 80°C for DNA extraction, for culture-based experiments soil was collected and preserved with 30% glycerol before storage at -80°C.

**Table 1:**
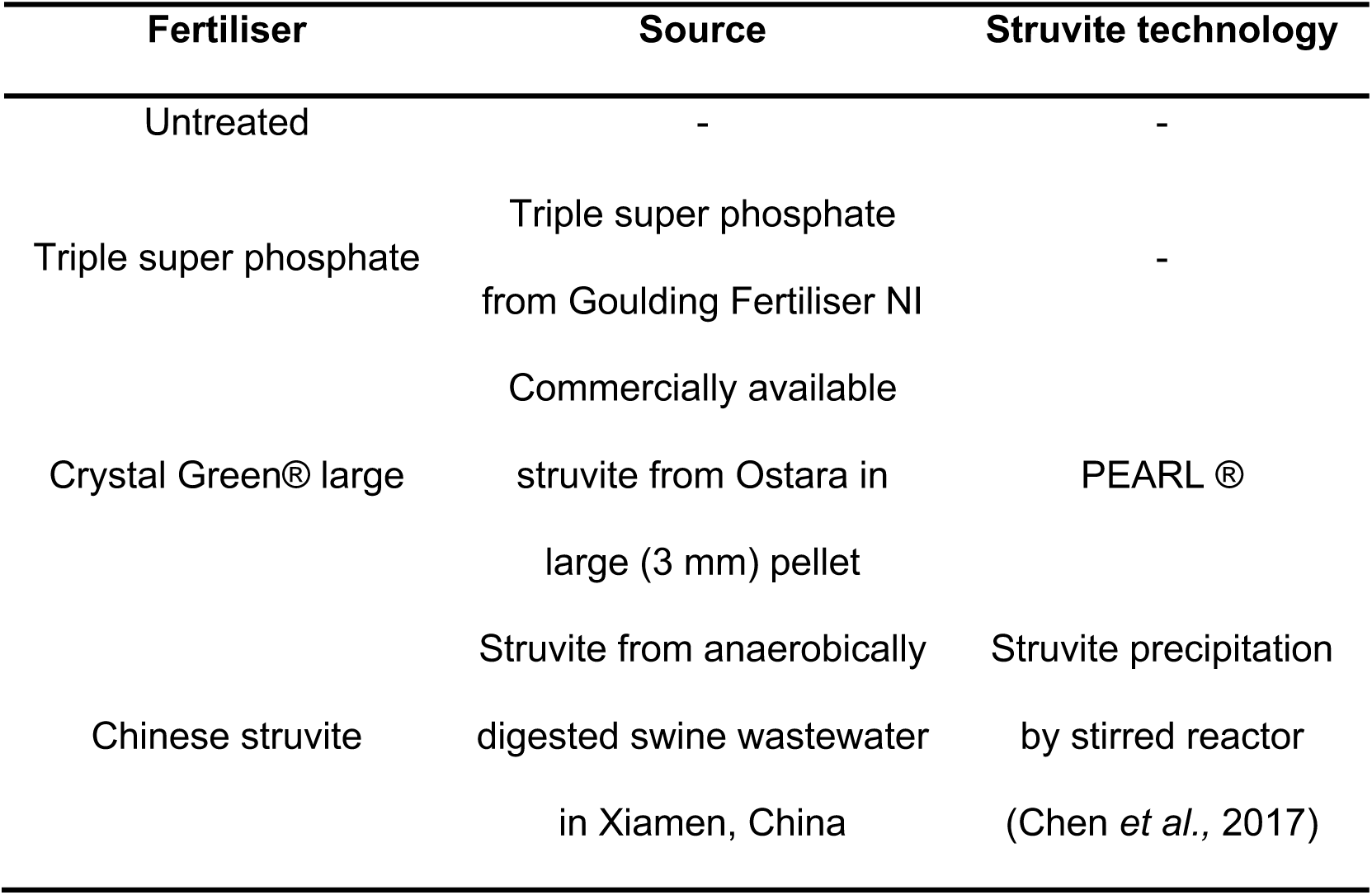
Summary of the fertilisers and treatments used by Hall *et al.,* 2020 and were also used in this study. Information adapted from Hall *et al.,*2020 Table 1.

### Antimicrobial residue analysis using liquid chromatography tandem mass spectrometry (LC-MS/MS)

Overall, > 130 pharmacologically active substances across 15 different drug classes, including antiparasitics and antibiotics, were included in the targeted LC-MS/MS method used for analysis of antimicrobial residues. Analysis of the samples was performed using LC-MS/MS following the method developed by Kenjeric *et al*. (2024). Briefly, a QTrap 5500+ MS/MS system (Sciex, Foster City, CA, USA) equipped with a Turbo V electrospray ionization (ESI) source was coupled to an ExionLC AD System (SCIEX). Chromatographic separation was performed at 25°C on a Gemini C18 column, 150 × 4.6 mm i.d., 5 μm particle size (Phenomenex, UK). Elution was carried out in binary gradient mode with a flow rate of 1 mL/min. Both mobile phases contained 5 mM ammonium acetate and were composed of methanol/water/acetic acid (10:89:1, v/v/v; eluent A) and (97:2:1, v/ v/v; eluent B) respectively. For further purification of reverse osmosis water, a Pure-lab Ultra system (ELGA Lab Water, Celle, Germany) was used. After an initial hold time of 2 min at 100 % A, the proportion of B was increased linearly to 50 % within 3 min. A further linear increase of B to 100 % within 9 min was followed by a hold time of 4 min at 100 % B, with the column then re-equilibrated at 100 % A for a further 2.5 min. The injection volume was set at 5 μL.

Extraction of the samples was performed as outlined in Kenjeric *et al*. (2024) (https://doi.org/10.1016/j.foodchem.2024.138834) with minor modifications. Briefly, a 1 g aliquot of sample was transferred to a 15 mL falcon tube and extracted with 4 mL of the extraction solvent [ACN:H20 (80:20, v/v)]. Samples were transferred to a multi-tube vortex shaker for 90 min, then centrifuged for 15 min at 5,000 rpm. A 500 μL aliquot of the supernatant was taken and diluted (1:1, v/v) with the same volume of diluent [ACN:H20 (20:80, v/v)], giving an overall dilution factor of 8-fold. As the method had been developed for milk and poultry feed, it was decided to conduct a trial extraction experiment on ‘blank’ soil (Mineral Gley) prior to analysis of the ‘real’ samples. This was done to ascertain the efficacy of the extraction protocol on soil, as it is known that some antibiotics are known to form chelates with metals, such as Mg^2+^, Ca^2+^, Fe^2+^, Cu^2+^ and Zn^2+^ (Khurana et al., 2021). The rationale was to see that of those substances spiked into the soil, which could be ‘recovered’ by the extraction. Therefore, 1 g aliquots of ‘blank’ soil were spiked with a working standard solution containing the pharmacologically active substances to achieve a concentration of 200 ng/g. The experiment was carried out in triplicate, with samples either spiked at this concentration and left overnight at room temperature, or for 1 hour before extracting as outlined above, before analysis using LC-MS/MS as outlined above.

### Resistome analysis using culture independent methods

#### DNA extraction, shotgun metagenomic sequencing, and bioinformatic analysis

To prepare for deoxyribonucleic acid (DNA) extraction, 500 mg of each soil sample was weighed and 978 µL of sodium phosphate buffer at pH 7.2 was added. DNA extractions were then performed using the standard protocol for the MPBio FastDNA™ SPIN Kit for soil (SKU:6560200-CF). The ZymoBIOMICS Microbial Community Standard (D6300) was included as a positive control and negative controls containing only kit reagents (kitome), were also included during the DNA extractions, and downstream analysis. DNA was quantified using Nanodrop (ThermoFisher Scientific, NanoDrop™ One/OneC Microvolume UV-Vis Spectrophotometer, ND-ONE-W), and samples diluted (if necessary) in TRIS 10 mM pH 8.0 to achieve a final DNA concentration of 8 ng/µL. DNA extraction was also performed, as described above, directly on 500 mg of Xiamen struvite fertiliser (in triplicate):no DNA could however be obtained, therefore sequencing and downstream analysis was not performed on this sample type.

DNA extracted from soil samples underwent library preparation and shotgun metagenomic sequencing at the QUB Genomic Core Technology Unit (GCTU). Paired end sequencing was performed using the Illumina NovaSeq 6000 S4 300 flow cell with a read length of 150 base pairs. FastQC files were produced and investigated for evidence of sequencing problems. All analyses described were completed on the QUB High Performance Cluster-Kelvin.

#### Metagenomic Analysis

Trimmomatic (v0.39, Bolger et al., 2014) was used to trim low quality bases, remove adapters and remove any reads without a corresponding pair. A ‘*sliding window’* approach was used to trim at any site where the average quality of any 4 sequential bases dropped below Q20. Finally, any remaining short reads (<50bp) after trimming were removed. To identify potential contamination within the samples, Bowtie2 (v2.5.1, Langmead, and Salzberg, 2012) was used to align the samples to the human genome (*GCF_000001405.40_GRCh38.p14*), using default parameters. MetaPhlAn (v4.0.6, Blanco-Míguez et al., 2023) was used to taxonomically assign the trimmed, filtered and decontaminated reads using default parameters. The MetaPhlAn reports were then used to calculate the taxonomic abundances at phylum, family and genus levels, and technical replicates averaged. The negative control was investigated for contamination, and the MetaPhlAn assigned taxonomy of the positive control compared to the expected Zymo mock community.

To identify ARGs within the cleaned metagenomic reads, two approaches were employed. DeepARG v1.0.2 was used, this is a deep learning-based approach to predict ARGs from metagenomes using their provided DeepARG-SS model (Arango-Argoty et al., 2018). The second approach used a read mapping protocol, which mapped the reads against two ARG databases using Bowtie2 (v2.5.1). The influence of bowtie2 parameters on the reported soil resistome were explored, including default (df, which reports the “best” alignments) and the -- very-sensitive-local alignment options, and with and without -a (which searches for and reports all alignments). We also investigated the effect of the chosen ARG database, comparing both the ResFinder database v2.4.0 (Bortolaia et al., 2020) and the CARD database v3.3.0 (Alcock et al., 2023), and how different post alignment filtering affected the reported resistome, including the minimum ARG coverage, and the minimum number of reads mapping to each ARG per sample to be considered as “present”. ARG coverage was calculated using SAMtools (v1.21) coverage, which calculates the coverage at each position; SAMtools defines coverage as the percentage of positions within each bin with at least one base aligned against it (Danecek et al., 2021). ARG abundances were then normalised as the number of reads mapped to the ARG per million of total reads per sample.

### Assessing the impact of Xiamen fertiliser on the culturable soil resistome

Xiamen struvite-treated soil and untreated soil (Time 0 hr and End point), which had been preserved in glycerol, were tested for ’culturable resistance’ against 7 antibiotics. Break points (BPs) were selected using all available Europeans Committee on Antimicrobial Susceptibility testing epidemiological cut-off values (EUCAST ECOFF) values (March 2024) for each antibiotic. These concentrations were then doubled to give the following project BPs: Amoxicillin (beta-lactam/penicillin, 32 mg/L), erythromycin (macrolides, 32 mg/L), florfenicol (synthetic phenicol, 32 mg/L), neomycin (aminoglycoside, 32 mg/L), tetracycline (tetracycline, 128 mg/L), trimethoprim (synthetic dihydrofolate reductase inhibitors, 32 mg/L) and streptomycin (aminoglycoside, 128 mg/L). Nutrient agar was autoclaved (121°C for 15 minutes) and cooled to 55°C before the addition of the antibiotic solutions, followed by plate pouring to achieve the final concentration of each BP. Nutrient agar without antibiotics was also prepared to determine the total bacterial count, including susceptible bacteria, for each sample. Stock solutions of soil were prepared by homogenising 270 mg of soil with 2430 µL of PBS, and tenfold dilutions were completed. One hundred microlitres of sample, ranging from the neat sample to 10^-3^ dilution, were plated in duplicate onto nutrient agar plates using a spread plate technique. The plates were then incubated aerobically at 25°C and colonies were counted after 24 hours and 48 hours, and colony forming unit per gram (CFU/g) calculated. Following 48 hours incubation, morphically distinct colonies were picked from plates inoculated with XIA Struvite treated soil (tE) and restreaked to purity from each antibiotic containing plate. These pure cultures were stored at -80 for further analysis.

### Taxonomic identification of AMR bacteria isolated from Xiamen struvite fertilised soil

Fifty-nine isolates obtained from the XIA fertilised soils were grown for 24-28 hours in Luria-Bertani (LB) broth, DNA was then extracted from 1 mL of broth using Powersoil beads for cell lysis and a MagNA pure 96 DNA kit (Rosch Diagnostic Limited). DNA was quantified using the Quant iT dsDNA high sensitivity assay. Negative controls were included during the extraction, PCR, and library preparation phases. Positive controls consisted of the Zymo mock community, which was included at the DNA extraction phase and in all downstream processing. The standard Nanopore 16S rDNA Barcoding Kit 1-24 (SQK-16S024) protocol was performed using the extracted DNA. Briefly this included amplification of the 16S rRNA gene using supplied barcodes, 10 μL of DNA was mixed with 5 μL of nuclease-free water and 25 μL of LongAmp Hot Start Taq 2X Master, 10 μL barcodes were then added. Due to the number of isolates, three barcode kits were used, and three separate libraries were prepared. The PCR cycle was as follows; initial denaturation at 95 °C for 1 min, then 25 cycles of denaturation at 95 °C for 20 secs, annealing at 55 °C for 30 secs, extension at 65 °C for 2 mins and a single cycle of final extension performed at 65 °C for 5 mins before being held at 4 °C. Following PCR the quality, size and integrity of the PCR products were assessed using D1000 ScreenTape and the TapeStation 4200 (Agilent Technologies). DNA was quantified using the Quant iT dsDNA high sensitivity assay previously mentioned. Libraries were then cleaned using AMPure XP beads. Each set of barcoded libraries were then pooled and rapid sequencing adapters added. Each DNA library was loaded onto a separate primed flow cell and sequenced on an Oxford Nanopore GridION. Taxonomic assignment was then performed, analysis began by trimming the adaptors using porechop v0.2.4 (default parameters, Wick, 2017), chopper v0.9.0 (De Coster and Rademakers., 2023) was then used to filter the reads within a minimum quality score of 10, a minimum length of 1250 and maximum length of 1700, finally EMU v3.5.0 (Curry et al., 2023) was used to assign taxonomy using default parameters and the EMU default database (Schoch et al., 2020). Taxonomic assignments at the genus level generated by EMU for the positive control samples were evaluated against the expected taxonomic distribution for the Zymo mock community. Negative controls were confirmed to show no detectable contamination by containing negligible DNA and reads.

### Statistics and data visualisation

All statistical analyses and figure preparation was performed in RStudio (v 2025.5.1.513, R v4.5.1). Diversity indices were calculated using the MetaPhlAn calculate_diversity.R scripts available from MetaPhlAn. Microbial community composition and alpha diversity were analysed in RStudio, using the packages vegan (v2.6.10, Dixon et al., 2003) and pairwiseAdonis (v0.4.1, Martinez Arbizu et al., 2020). Bray-Curtis dissimilarities were calculated on log-transformed genus- and phylum-level normalised read data, and permutational multivariate analysis of variance (PERMANOVA) with 999 permutations was used to investigate the effects of treatment and time. Pairwise PERMANOVA comparisons were performed across all relevant experimental groups, with p-values adjusted for multiple comparisons using the Benjamini-Hochberg method. The MetaPhlAn computed Jaccard and Bray-Curtis distance matrices were also analysed to validate community composition differences. Alpha diversity, measured by Simpson and Shannon indices, was assessed using Wilcoxon rank-sum and pairwise Wilcoxon tests, with Benjamini-Hochberg correction applied. All results were exported to Excel for downstream reporting.

Normalised ARG abundance data were analysed using linear mixed-effects models with the lmer() function from the lme4 package (v1.1.37). For each resistance gene, a separate model was fitted with treatment, timepoint, and their interaction (Treatment * Time) as fixed effects, and technical replicate as a random effect to account for repeated measurements. Post hoc pairwise comparisons between treatment groups at each timepoint were conducted using the emmeans package (v1.11.1), with estimated marginal means and contrasts. The resulting p-values from all gene-wise comparisons were corrected for multiple testing using the Benjamini-Hochberg procedure to control the false discovery rate (FDR).

Pairwise T tests to compare log transformed CFU/g data against treatment and time point were completed using base R, the data was log transformed, tested and the pvalues adjusted using the Bonferroni method. Permutational multivariate analysis of variance (PERMANOVA) was used to assess the effect of treatment and time on the metagenomic taxonomic data and was conducted using adonis() function in the vegan package, employing 999 iterations. The representative genus level tree was generated using the NCBI Common Tree tool. The output tree was reformatted with PHYLIP (v3.695, Felsentein et al., 1993) in preparation for visualisation and annotation using ggtree (v3.16.3, Yu et al., 2017).

## Results

### Antimicrobial residues as identified by LC-MS/MS

LC-MS/MS was used to assess the presence of antimicrobial compounds in both fertiliser-only and soil-fertiliser mixtures. The Xiamen (XIA) struvite fertiliser was found to contain the antimicrobial compounds monensin, nicarbazin, pipemidic acid, and sulfadoxine at varying concentrations (Table 2). Notably, XIA_1 (technical rep1) had higher levels of monensin, pipemidic acid, and sulfadoxine, with the latter two compounds not detected in other replicates. In contrast, XIA_3 exhibited high levels of nicarbazin, whereas the other two replicates contained only trace amounts, indicating variability in antimicrobial compound composition. In comparison, TSP and CG fertilisers contained no detectable antimicrobial compounds beyond trace levels (< 1 μg/kg). Similarly, soil fertilised with XIA, TSP, and CG only showed trace amounts of closantel and thiabendazole [Supplementary table 1]. One caveat on the analytes detected, is that based on the experiment conducted on the ‘blank’ soil to determine which antimicrobials could be successfully extracted, 14% of those spiked could not be recovered after an o/n spike, whereas another 21% had recoveries of extraction of < 50% [Supplementary table 2]. Therefore, there could be more analytes present than detected, possibly due to metal chelation and therefore being non-extractable as indicated by the recovery experiment.

**Table 2:**
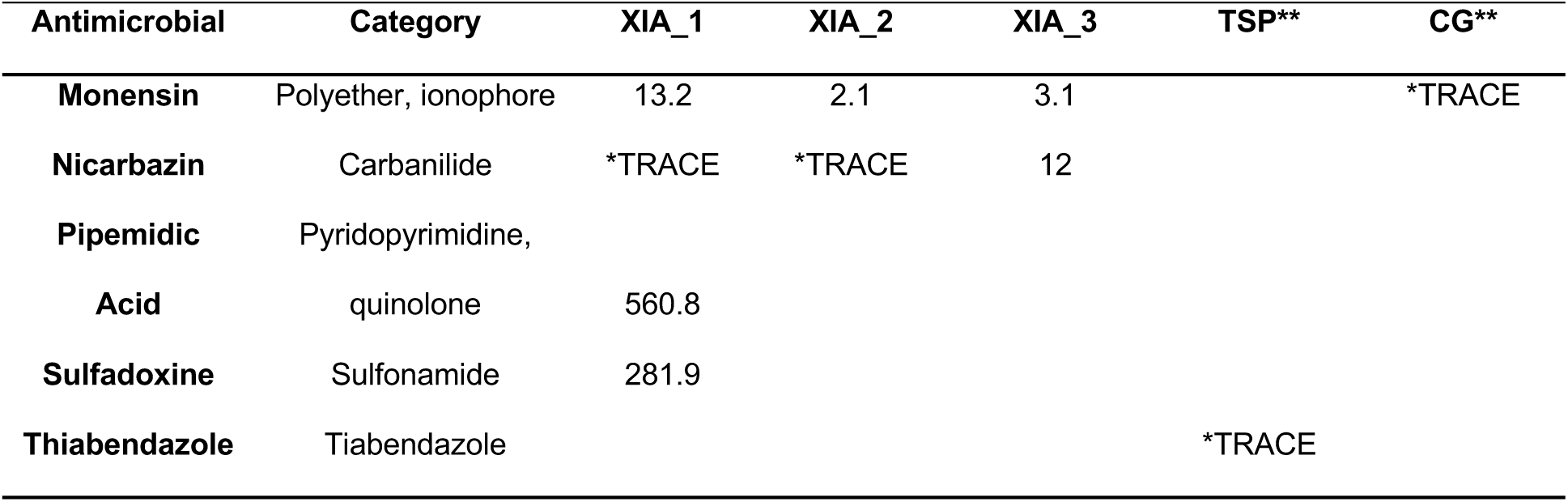
LC-MS/MS analysis of fertiliser samples to identify antimicrobial compounds, concentrations (μg/kg) of detected compounds are shown. “*TRACE” indicates compounds detected at trace levels below the quantification limit; less than lowest calibrant of 1 μg/kg (1 ppb), ** 3 replicates analysed and average. Note 1, 2, and 3 represent technical reps.

| Antimicrobial | Category | XIA_1 | XIA_2 | XIA_3 | TSP** | CG** |
| --- | --- | --- | --- | --- | --- | --- |
| **Monensin** | Polyether, ionophore | 13.2 | 2.1 | 3.1 |  | *TRACE |
| **Nicarbazin** | Carbanilide | *TRACE | *TRACE | 12 |  |  |
| **Pipemidic Acid** | Pyridopyrimidine, quinolone | 560.8 |  |  |  |  |
| **Sulfadoxine** | Sulfonamide | 281.9 |  |  |  |  |
| **Thiabendazole** | Tiabendazole |  |  |  | *TRACE |  |

### Fertiliser conditions do not induce significant taxonomic shifts

Twenty-four soil samples, representing three different fertiliser amendments and a control condition at two time points (start and end), underwent shotgun metagenomic sequencing. The average number of reads per sample was 7.53 × 10⁹. The negative control contained 11,788 raw reads; following the read mapping decontamination step, only 720 reads remained, 100% of which could not be classified by MetaPhlAn, while the positive control was comparable to the expected taxonomy of the Zymo mock community [Supplementary table 3] (25,992,483 paired end (PE) raw reads, 23,925,235 PE reads following trimming and decontamination). After quality control (cleaning and trimming), the average number of reads across soil samples was 44,783,192 [full taxonomic classification is available in Supplementary Table 4].

Taxonomic classification by MetaPhlAn revealed that microbial composition across soil samples was highly similar, with no significant differences between treatments at either the phylum or genus level (p.adj ≥ 0.05, Supplementary Table 5), suggesting there were no observable impacts of CG, XIA, and TSP treatment on the bacterial and archaeal diversity in the soil. At the kingdom level, soil samples contained an average of 95.23% Bacteria and 4.76% Archaea, and 0% of reads were unable to be classified across samples. A total of 14 different phyla were identified, of which eight were present in all samples, while the remaining six were found at low levels, with an average relative abundance of 0.114% across samples [Supplementary Figure 1]. The most abundant phyla were Proteobacteria (31.58%), Verrucomicrobia (23.15%), and Actinobacteria (22.32%). Of the 14 identified phyla, 11 belonged to the kingdom Bacteria, while three were from the kingdom Archaea: Thaumarchaeota, Euryarchaeota, and Candidatus Thermoplasmatota. The archaeal phyla were generally found at lower abundances, with average relative abundances of 4.745%, 0.013%, and 0.001%, respectively, across the 24 samples.

At the genus level, a total of 117 different genera were identified across the soil metagenomes, with average relative abundances ranging from 22.72% to 0.00423%. On average at genus level, 7.62% of reads were unable to be classified. Twenty genera were present in all samples. The most abundant genera included *GGB32810* (p Verrucomicrobia), *GGB61943* (p Proteobacteria), *GGB52803* (p Actinobacteria), *GGB66002* (p Proteobacteria), and an unclassified genus belonging to the family *Acidobacteriaceae* (p Acidobacteria), with average abundances ranging from 22.72% to 4.92% (Figure 1). There were no significant changes in microbial diversity due to treatment, as assessed by Simpson, Shannon, Jaccard and Bray-Curtis analyses (p.adj ≥ 0.05) [Supplementary Table 5].

**Figure 1:**
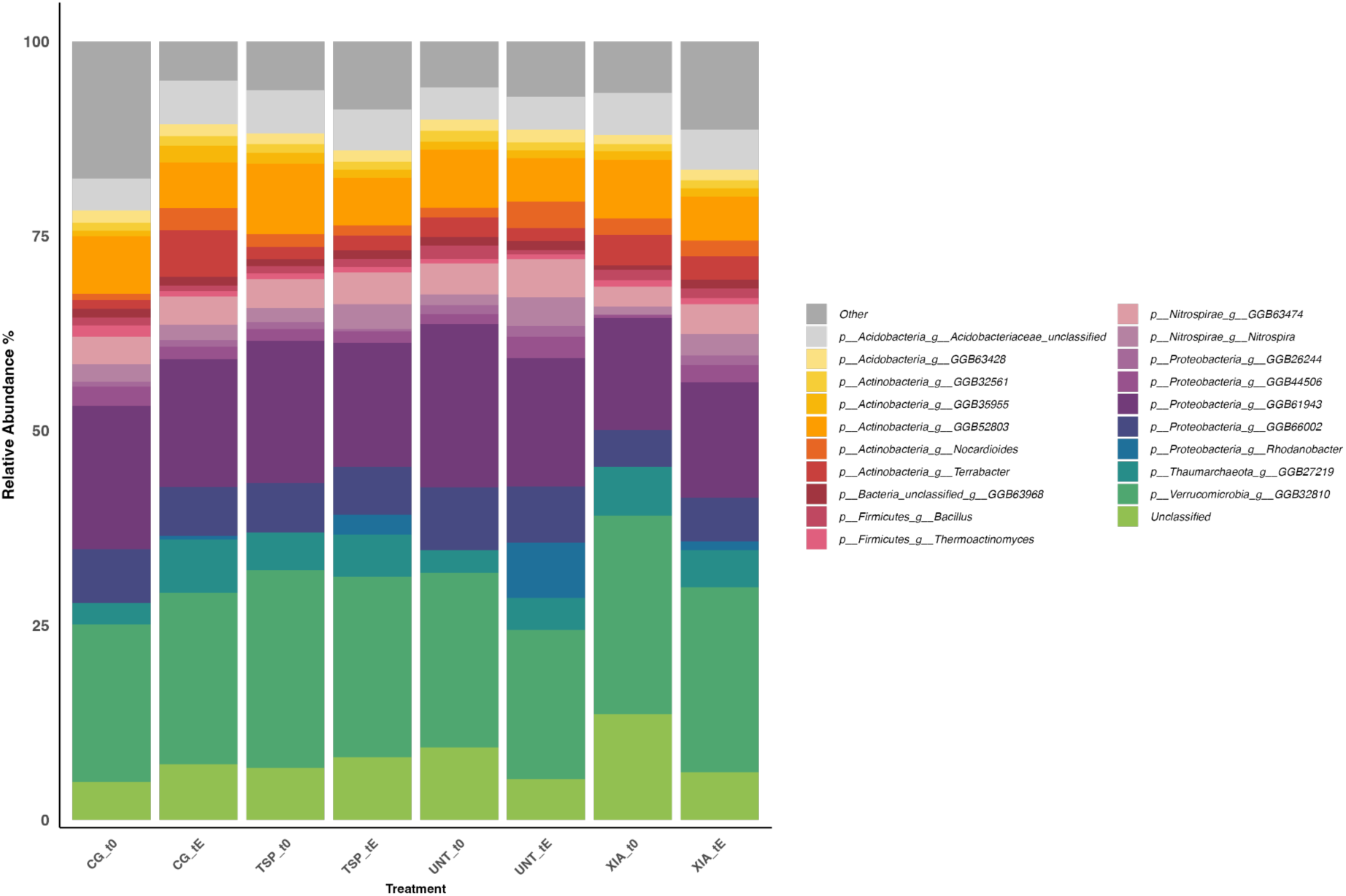
Taxonomic composition of soil metagenomic reads assigned using MetaPhlAn. Stacked bar plots display the relative abundance of the top 20 most abundant genera across treatments and points (t0 = time zero, tE = end time point), all other genera grouped as "Other. UNT (Untreated), CG (Crystal Green), XIA (Xiamen), and TSP (Triple Super Phosphate).

### Tool, database, and parameter choices greatly affect the resistome profiles observed in soil metagenomes

The curation and availability of ARG databases, such as those provided by CARD and ResFinder, have driven the development of tools to identify ARGs in sequence data. Among these, DeepARG is one of the most widely used, with around 800 citations since its publication in 2018, and it remains popular today. DeepARG combines deep learning with more traditional sequence alignment methods (such as DIAMOND (Buchfink et al., 2021)) to improve ARG predictions and reduce false negatives by “…not requiring strict cutoffs, which enables identification of a much broader diversity of ARGs.” However, although DeepARG reported a high number of ARGs in our samples, closer inspection revealed that average gene coverage was low (24.1%), with the majority of ARGs having alignments that covered less than 17.5% of the reference gene [Supplementary table 6]. Only a small fraction (2.9%) of detected ARGs had coverage exceeding 90% and in some cases, ARG presence reported by DeepARG was based on a single mapped read [Supplementary table 6].

Using a single sample of results to highlight some of these issues and confusion regarding outputs (PN0204_0336_S330, UNT_t0_Rep1), DeepARG (run with default parameters) reported an average ARG coverage of just 25.8%. For example, *AAC(3)-I* was detected based on only a single read with an alignment length of 34 amino acids, despite the reference gene being 177 amino acids long. In total, 154 ARGs were identified in this sample, 39 of which were supported by only a single read, and 128 ARGs by fewer than 100 reads. Even ARGs with relatively high read counts showed limited coverage. *OTRA* had 107 mapped reads, all aligning to positions 1–144 of a 663 amino acid gene, giving just 21.7% coverage. *MUXC* (1036) had 322 reads, but these primarily aligned to hotspots, yielding a total coverage of only 37.9%. In contrast, *RPHB* had 364 aligned reads spanning multiple regions across its 884 amino acid length, resulting in 93.4% coverage; this ARG was also identified using our alternative mapping-based approach (described below).

In an attempt to reduce the number of low-coverage and thus spurious ARGs reported, we applied the --gene_coverage parameter in DeepARG, which sets the minimum percentage of gene coverage required to consider a full gene. However, modifying this parameter to 0.2 or 0.9 (from the default of 1) resulted in no change to the reported ARG profiles, raising questions about how this setting functions in practice and further highlighting issues related to low-coverage ARG calls and output inconsistencies [Supplementary table 7].

To address the concerns identified with DeepARG’s output, we next explored a more direct and transparent approach: read mapping using Bowtie2. This strategy was chosen to provide greater confidence and interpretability in the results by relying on a well-understood alignment process, where both the procedure and the meaning of the output are more easily verified. Overall, more ARGs were identified using the CARD database than ResFinder (Figure 2). Both databases yielded low ARG counts under the default bowtie2 alignment parameters, regardless of the -a setting (which reports all alignments), with a maximum of three ARGs detected across the 24 samples (when no read threshold was applied). When minimum read thresholds (e.g., 50 or 100 reads) were introduced to the default bowtie2 alignment, 0 ARGs were detected across all samples, regardless of database or -a usage (Figure 2). In contrast, the -very-sensitive-local (--vsl) alignment setting detected a higher number of ARGs overall (Figure 2).

**Figure 2.**
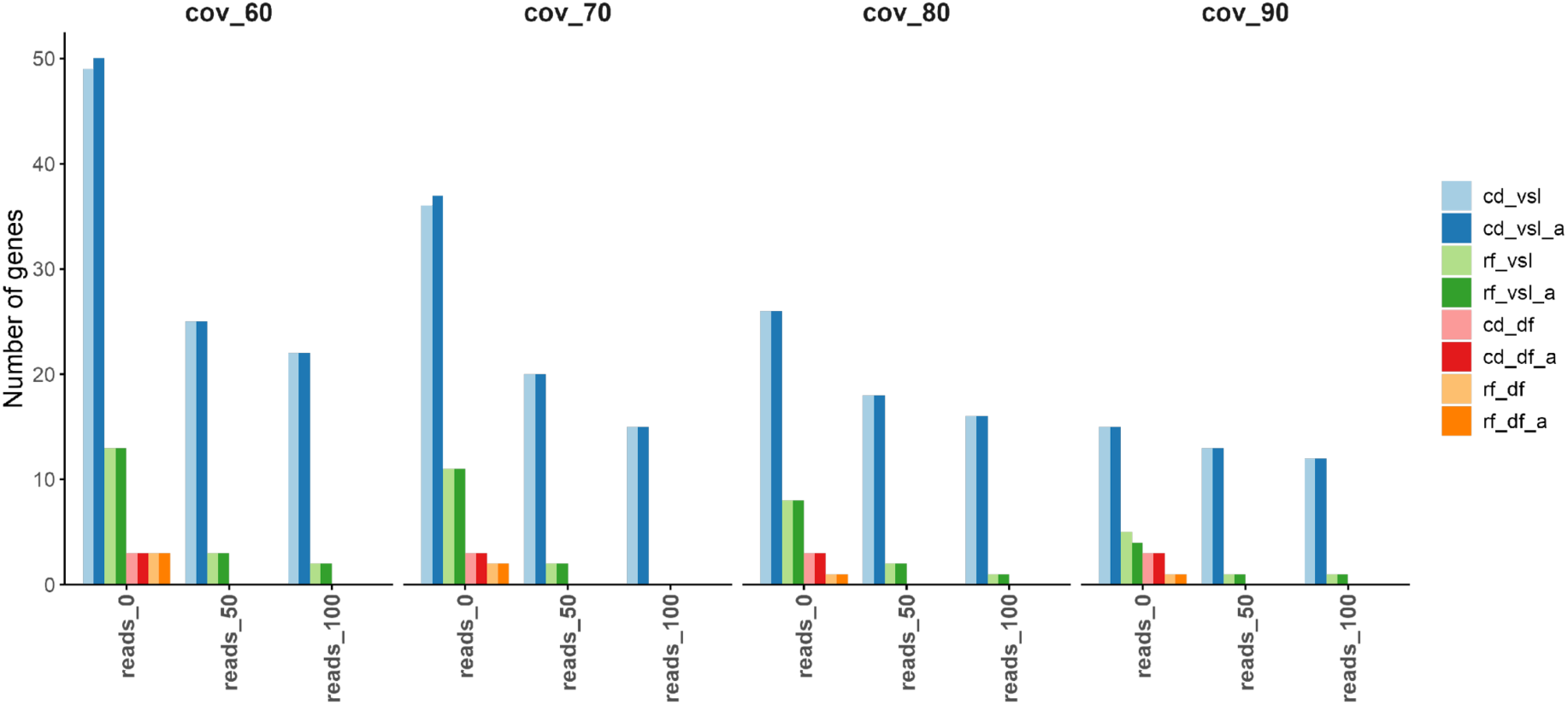
Number of unique antimicrobial resistance genes (ARGs) detected from soil metagenomes under varying analytical parameters. Comparisons were made across the following: **(i)** databases-CARD (RGI-based; *cd*) and ResFinder (*rf*); **(ii)** read mapping parameters-Bowtie2 with default settings (*df*) versus *--very-sensitive-local* (*vsl*), with the ‘-a’ flag (report all alignments) either enabled or disabled; and **(iii)** detection thresholds-minimum number of mapped reads (0, 50, or 100) and minimum gene coverage cutoffs (60%, 70%, 80%, or 90%).

Increasing the minimum coverage threshold led to a substantial reduction in detected ARGs. For example, CARD (with -a), ARG counts dropped from 50 to 15 when the coverage threshold increased from 60% to 90%; for ResFinder, they decreased from 13 to 4 (Figure 2). At lower coverage thresholds (60% and 70%), applying minimum read filters dramatically reduced ARG detection. For example, at 60% coverage, CARD’s ARG count dropped from 50 to 22, and ResFinder’s dropped from 13 to 2 (with a 100-read minimum). At 70% coverage, ResFinder’s ARGs dropped from 11 to 0 (Figure 2). At higher coverage thresholds (e.g., 90%), the impact of read filtering was less pronounced, with CARD ARG counts remaining stable (12–15), regardless of read thresholds (Figure 2).

Finally, we assessed the impact of the -a option (reports all alignments) on read mapping under the --vsl alignment, 90% gene coverage, and a 100-read minimum threshold. For ResFinder, only one ARG (*ole(C)_1_L06249*) was detected, with 3,339 reads mapped without -a and 3,351 with -a, across five samples it was detected in, an increase of just 12 reads (average per sample: 667.8 vs. 670.2) [Supplementary table 8]. Of the 12 CARD-identified ARGs, eight showed minimal differences (<20 reads), while 4 showed more substantial increases with -a: *Sven_rox* (+214), *MuxB* (+2,193), *ceoB* (+2,622), and *rpoB2* (+10,559). For *rpoB2*, although it was detected in the same 18 samples with or without -a, the average read count per sample was 586.6 higher with -a.

Therefore, the following parameters were selected for downstream analysis: -a, --very-sensitive-local (--vsl) alignment, 90% coverage threshold, and a minimum of 100 mapped reads per ARG. By using a stringent 90% coverage threshold, we aim to minimize false positives caused by reads mapping to widely shared conserved domains rather than the full ARG and limiting the risk of overestimating gene presence. Using ResFinder under these conditions identified a single ARG, *ole(C)_1_L06249*, which was also detected using the CARD database. This ARG appeared in the same five samples across both databases, with nearly identical read counts, only one sample differed by two reads (751 vs. 749) [Supplementary table 7]. Consequently, downstream results are presented using the CARD data, with ARG counts normalised to reads per million per sample (Figure 3).

**Figure 3:**
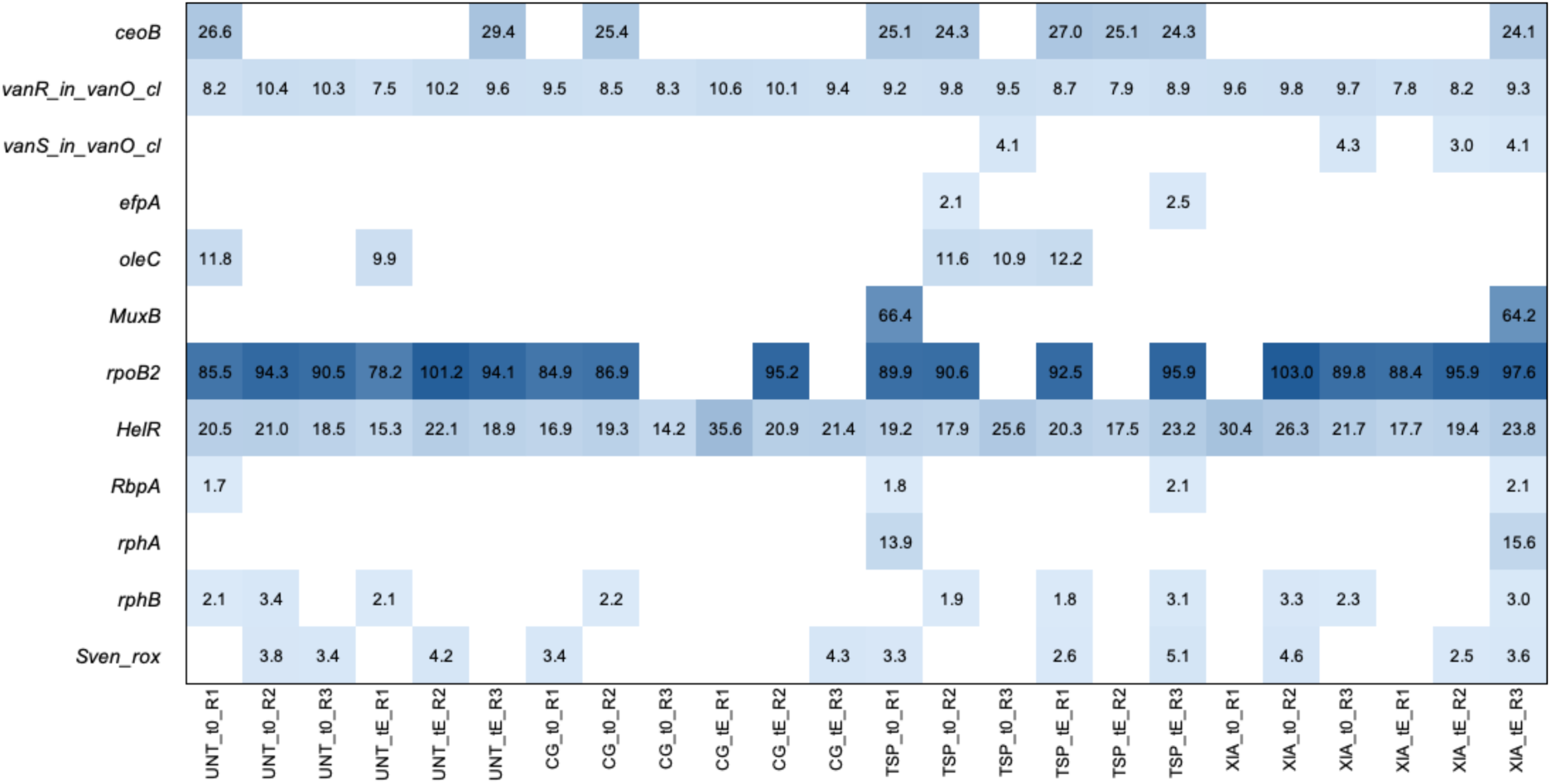
Heatmap showing normalised abundance of antibiotic resistance genes (ARGs) detected in metagenomic samples across treatments and timepoints. Each cell represents a single replicate. ARG abundance is expressed as the number of reads mapped per million total reads. ARGs were identified using the CARD database with Bowtie2 (--very-sensitive-local) alignment. A 90% gene coverage threshold and a minimum of 100 mapped reads per ARG were applied for inclusion. Darker shades of blue indicate higher ARG abundance.

### Treatment and time did not significantly affect ARG abundance in soil

Across all timepoints and treatments, a total of 12 ARGs were identified, with all 24 soil samples containing at least two ARGs (Figure 3). These genes conferred resistance to the following antibiotic classes (number of ARGs in parentheses): aminoglycoside + fluoroquinolone (1), glycopeptide (2), isoniazid-like + rifamycin (1), macrolide (1), multidrug resistance (1), peptide + rifamycin (1), and rifamycin (5). The most widespread ARGs were *vanR_in_vanO_cl* (glycopeptide), *rpoB2* (peptide + rifamycin), *HelR*, and *Sven_rox* (both rifamycin), which were detected in at least one replicate across all treatments and timepoints. Notably, *vanR_in_vanO_cl* and *HelR* were present in every sample. While the genes with the highest abundance across samples were *rpoB2* (peptide + rifamycin) and *HelR* (rifamycin) (Figure 3).

Overall, fluctuations in ARG abundance were small, with many genes exhibiting a decline in abundance over time (Figure 3). The largest increase within a treatment was observed for *rpoB2* in XIA-treated soil, where its average abundance rose from 64.3 at t0 to 94.0 at tE. In contrast, the same gene showed the largest decrease, declining from 57.2 to 31.7, a 44.6% reduction in CG treated soil. Importantly, no ARGs were significantly different in abundance due to treatment or time (p.adj ≥ 0.05).

### XIA struvite fertilisation significantly increased the total bacterial load and culturable AMR bacteria

To assess the presence of AMR bacteria, soil samples were inoculated onto agar plates containing various antibiotics. Following incubation, bacterial colonies were isolated, and DNA was extracted for taxonomic identification using 16S rRNA gene sequencing. No bacterial growth was observed on agar plates containing tetracycline-supplemented agar inoculated with soil. For erythromycin, florfenicol, and neomycin, only low colony growth was detected, with fewer than 30 colonies present, making CFU/g calculations unreliable after 48 hours. However, despite the inability to quantify CFU/g, colonies were successfully isolated, purified, DNA-extracted, and 16S rRNA gene sequenced for downstream analysis. In contrast, CFU/g values were reliably calculated for amoxicillin, trimethoprim, and streptomycin at both 24-hour and 48-hour time points, along with the antibiotic-free control and are presented in Figure 4.

**Figure 4:**
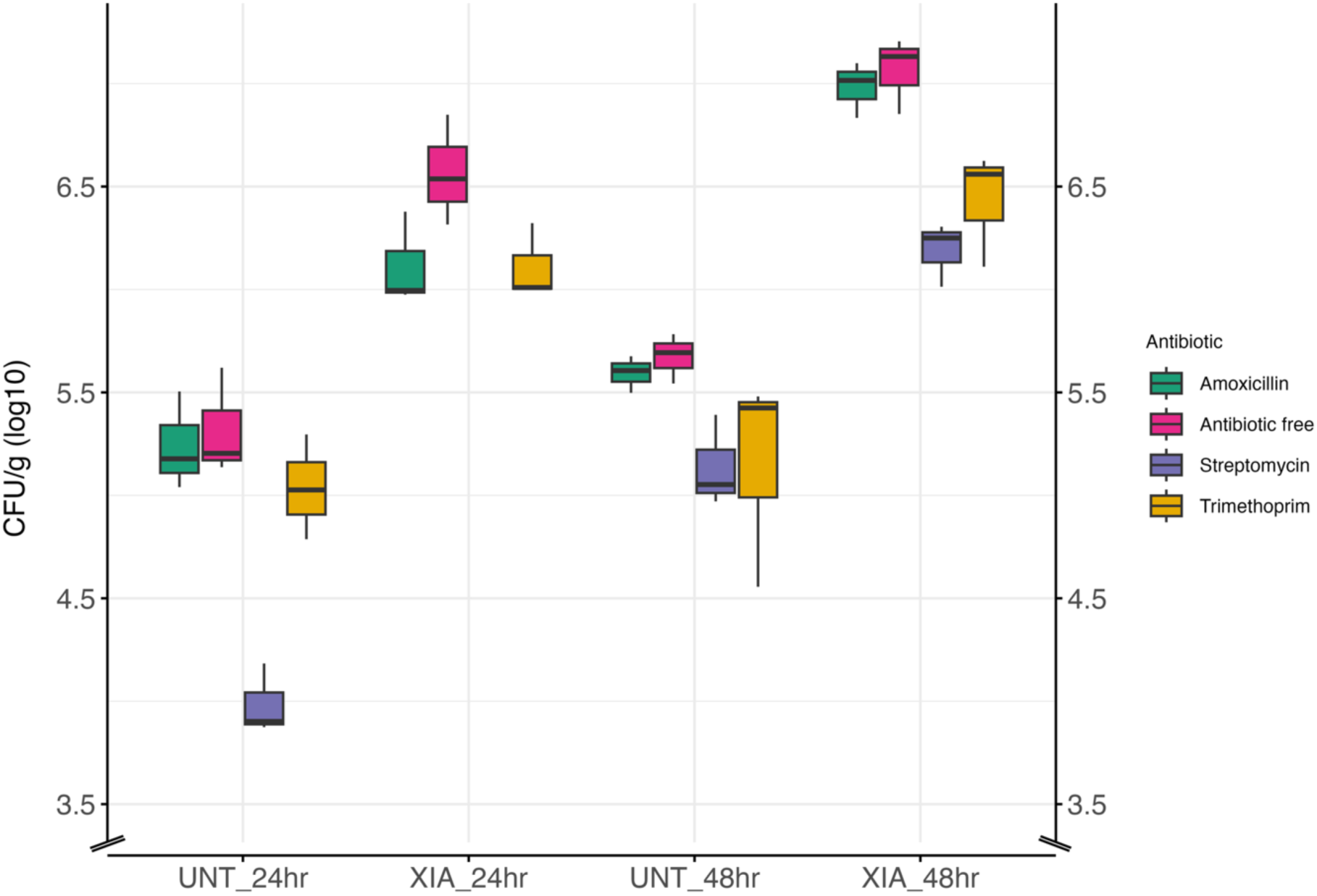
Boxplot of log₁₀-transformed CFU/g of soil across experimental groups, showing the effect of treatment and timepoint. Each box represents the distribution of three technical replicates, grouped by condition and coloured by antibiotic type: Amoxicillin (green), Trimethoprim (yellow), Streptomycin (purple), and Antibiotic-free (pink). Boxes display the median and interquartile range (IQR), with whiskers extending to 1.5×IQR. Note, the y-axis includes a break between 0 and 3.5 to accommodate the data range.

**Figure 5:**
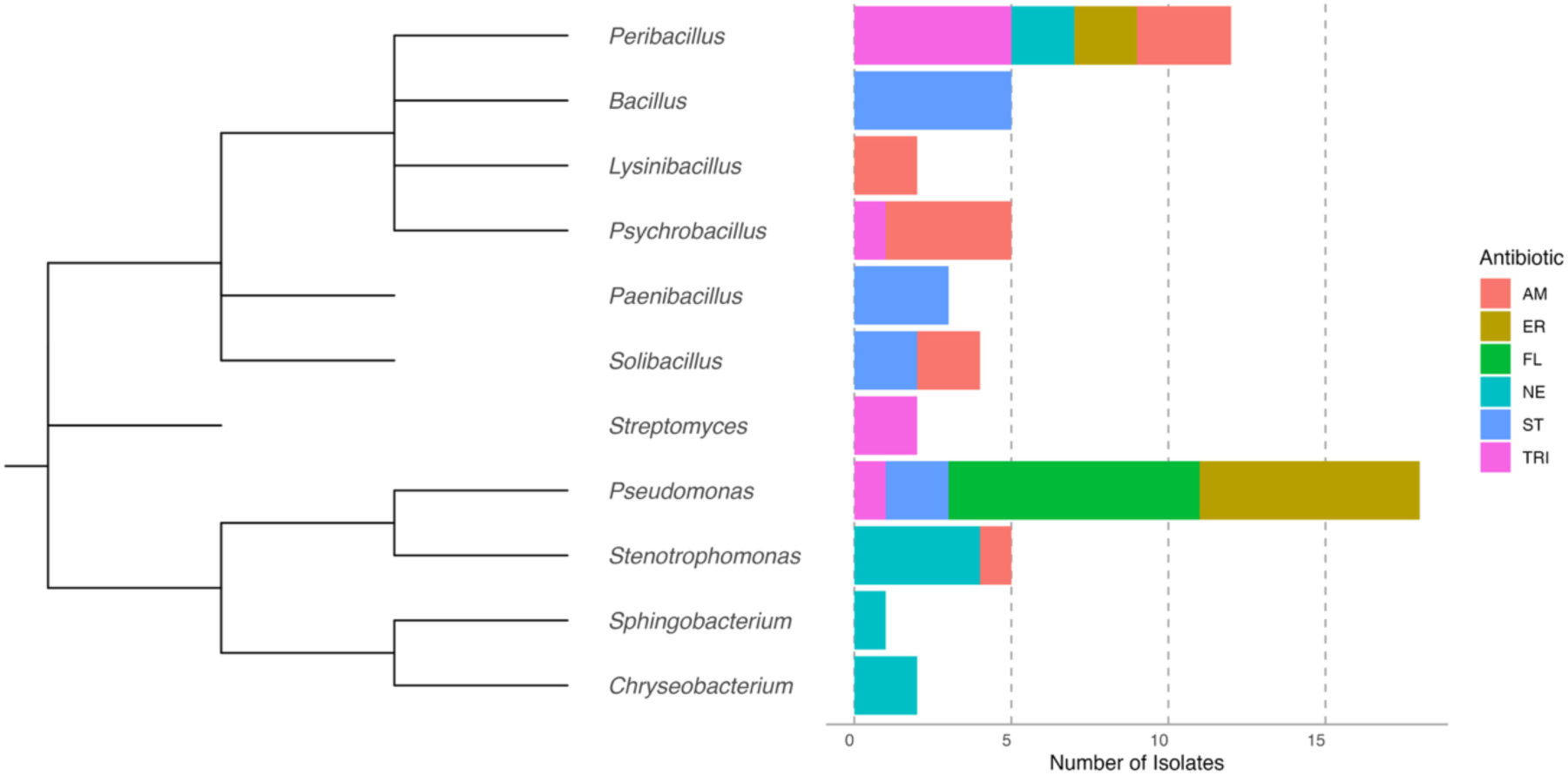
Phylogenetic distribution and antibiotic resistance profiles of AMR isolates from XIA-fertilised soil. A genus level representative tree shows relationships between 11 genera of AMR bacteria isolated from XIA-treated soil. The stacked bar chart (right) represents the number of isolates per genus and the specific antibiotics to which they were resistant to. AM (Amoxicillin), ER (Erythromycin), FL (Florfenicol), NE (Neomycin), ST (Streptomycin), and TRI (Trimethoprim).

**Figure 6:**
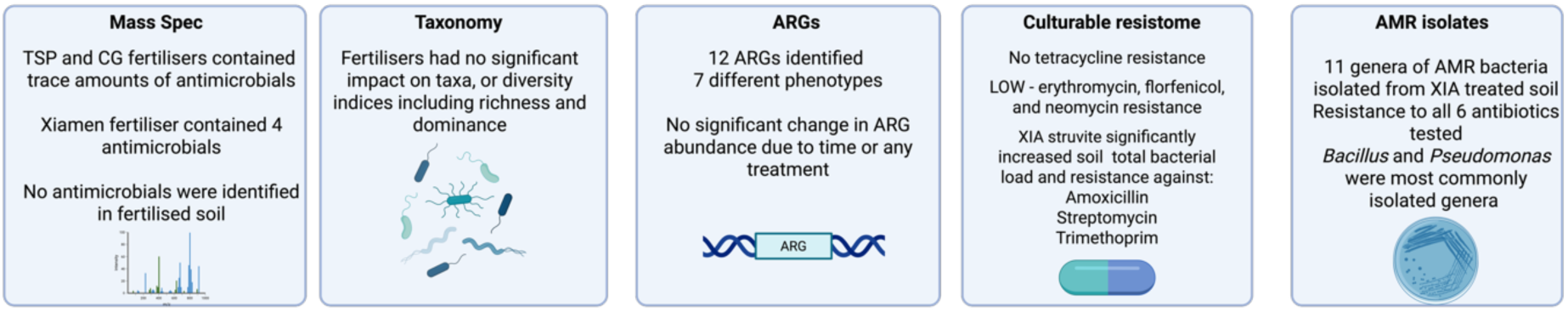
Summary of key findings on antimicrobial resistance (AMR) in fertilised soils. Five panels summarise major results across different analyses: mass spectrometry, taxonomic analysis (soil metagenomic reads), ARG profiling (soil metagenomic reads), culturable resistome analysis and AMR bacterial isolates from XIA fertilised soil.

XIA-treated soils consistently exhibited higher CFU/g values compared to untreated soils across all tested conditions and timepoints (Figure 4). The highest CFU/g values for amoxicillin, trimethoprim, streptomycin, and the antibiotic-free control were all observed in XIA-treated soil at 48 hours, with average CFU/g of 9.93x10^6^, 3.12x10^6^, 1.62x10^6^ and 1.23x10^7^ respectively. Direct comparisons between UNT and XIA-treated soils at both 24 and 48 hours showed significantly higher CFU/g values in XIA-treated samples for every condition tested (p.adj < 0.05), suggesting that XIA fertiliser promotes overall bacterial growth (Figure 4).

Within each treatment group, antibiotic-specific effects were also observed (Figure 4). At 24 hours, no significant differences were seen between amoxicillin, trimethoprim, and the antibiotic-free control in either untreated or XIA-treated soils (p.adj ≥ 0.05). However, streptomycin treatment led to significantly lower log CFU/g values compared to all other conditions in both soil types (p.adj < 0.05), indicating stronger suppression of bacterial growth (Figure 4). By 48 hours, antibiotic effects diverged further. In UNT soil, no significant differences were observed among any treatment conditions (p.adj ≥ 0.05). In contrast, XIA-treated soil still showed significantly lower log CFU/g under streptomycin compared to amoxicillin (p.adj = 0.026) and the antibiotic-free condition (p.adj = 0.008) (Figure 4).

Temporal comparisons (48 hr vs. 24 hr) revealed minimal changes in UNT soils, except for streptomycin, which showed a significant increase in log CFU/g over time (p.adj = 0.0001). In XIA-treated soil, log CFU/g values significantly increased over time under both amoxicillin (p.adj = 0.009) and streptomycin (p.adj = 5.52 × 10⁻²⁴) treatments, while trimethoprim and the control remained stable (Figure 4). Overall, streptomycin consistently suppressed bacterial growth more effectively than the other antibiotics at 24 hours but reached comparable CFU/g levels by 48 hours, particularly in XIA-treated soils (Figure 4).

### Diverse bacterial isolates in Xiamen fertilised soil exhibit resistance to six antibiotic classes

Taxonomic assignments generated by EMU for the positive control samples were evaluated against the expected taxonomic distribution of the Zymo mock community. EMU successfully identified the correct taxa in the positive control (ZYMO mock community standard), though the proportions differed from the expected distribution [Supplementary table 9]. Negative controls contained 5, 4,18 and 14 reads, confirming no detectable contamination, as they exhibited negligible DNA and read counts.

A total of 59 isolates were cultured from XIA-treated soil samples, exhibiting resistance to six tested antibiotics and belonging to 11 different genera. The predominant resistant bacteria were *Pseudomonas* (18 isolates) and *Bacillus* (12 isolates). *Pseudomonas* isolates displayed resistance to erythromycin (7 isolates), florfenicol (8), streptomycin (2), and trimethoprim (1), while *Bacillus* isolates exhibited resistance to amoxicillin (3 isolates), erythromycin (2), neomycin (2), and trimethoprim (5). Notably, *Bacillus* and *Pseudomonas* were the only genera containing isolates resistant to four antibiotic classes. This may be due to a range of reasons including intrinsic resistance mechanisms, horizontal gene transfer, or adaptive responses that allow these genera to persist in antibiotic-exposed environments. In contrast, three other genera, *Peribacillus*, *Solibacillus*, and *Stenotrophomonas*, contained isolates resistant to two antibiotic classes, while the remaining six genera harboured isolates resistant to only one class. In summary, a diverse range of drug-resistant taxa were able to be cultured from XIA fertilised soil.

## Discussion

P is an essential element for plant growth and a key resource in modern food production systems (Cordell and White, 2011; Ågren *et al.,* 2012), yet easily accessible, high quality, phosphorus rock resources are declining. Struvite is considered as a sustainable P containing fertiliser and could help to close the broken P cycle (European Commission, 2018). However, the impact of struvite-based fertilisers on the soil resistome remains unclear. Therefore, the AMR associated risk of struvite fertilisers must be carefully considered. In this study we evaluated the impact of Xiamen struvite fertiliser, produced from swine wastewater, on the soil resistome of rhizospheric soil planted with ryegrass following a 3-month growth period. Furthermore, we compared Xiamen struvite produced from piggery wastewater (XIA), which is not commercial grade, to a commercially available struvite, Crystal Green (CG), and to the commonly used inorganic phosphate fertiliser Triple Super Phosphate (TSP).

### No evident correlation between antibiotic residues in struvite fertiliser and soil ARG abundance

The commercially available fertilisers Crystal Green struvite and Triple Super Phosphate both contained only trace amounts of antimicrobials. However, the antibiotic concentrations found in XIA struvite were more variable. Although the XIA struvite did contain antibiotics, little to no antibiotic residue was identified in the soil (only trace amounts). This suggests that either the antibiotics are not being transferred, only small amounts are, or they are undergoing chelation and are therefore no longer detectable. When compared to XIA struvite used in other studies (Chen et al., 2017), the XIA fertiliser used here had generally lower antibiotic content, highlighting the variability of non-commercial struvite production.

Pipemidic acid, a fluoroquinolone, was detected at high levels in one replicate of XIA struvite. A fluoroquinolone resistance gene, *ceoB*, which encodes a cytoplasmic membrane component of the CeoAB-OpcM efflux pump, (Guglierame et al., 2006; Somprasong et al., 2021), was also found here in soil fertilised with XIA struvite. However, our current understanding is that to produce this efflux pump, and therefore a resistant phenotype, *ceoB* must be contained in an operon with the ARGs *ceoA* and *opcM* (Guglierame et al., 2006), neither *ceoA* nor *opcM* which were detected in this study. Additionally, *ceoB* was also present in untreated and non-struvite-treated soils, suggesting it is not specifically linked to struvite fertilisation. This antibiotic, pipemidic acid, is currently not approved for use in the EU in humans, due to the disabling and potentially permanent side effects caused by quinolone and fluoroquinolone antibiotics (European Medicines Agency, 2019). Similarly, a sulfonamide antibiotic was identified in XIA fertiliser, and within XIA fertilised soil the resistance gene *MuxB* was identified, which can confer resistance to sulfonamides. However, MuxB is only one of the two required components in the *Pseudomonas aeruginosa* efflux pump system MuxABC-OpmB (Mima et al., 2009).

These above-mentioned genes, *MuxB*, *ceoB*, and *efpA,* are associated with multidrug efflux mechanisms and are broadly distributed among soil-dwelling microbes such as *Mycobacterium* and *Xenophilus* spp. Additionally, soil-dwelling *Mycobacterium* species such as *Mycobacterium obuense* and *Mycobacterium rhodesiae* have been identified as hosts of *efpA*, as well as rifamycin and aminocoumarin resistance genes, highlighting the role of environmental mycobacteria as important reservoirs and vectors for ARG dissemination (Yang et al., 2022). *EfpA*, an MFS-type efflux pump originally characterised in *Mycobacterium tuberculosis*, is homologous to members of the *QacA* transporter family and contributes to resistance against multiple antibiotics, including isoniazid, fluoroquinolones, and rifamycin (Doran et al., 1997; Hashemzadeh et al., 2025). Studies have shown *efpA* to be significantly enriched, up to fourfold, in microplastic-rich plastisphere environments (Yang et al., 2022).

Among the ARGs identified, rifamycin resistance, represented by genes such as *efpA*, was the most prevalent phenotypic class. Firstly, this may be because rifamycin is naturally produced by soil-dwelling organisms. It was originally identified as a product of *Amycolatopsis rifamycinica* (formerly *Streptomyces mediterranei*), a soil-associated bacterium, and members of the *Amycolatopsis* genus are widely distributed in soil (Kisil et al., 2021). Secondly, rifamycin resistance has been reported to carry no significant fitness cost in soil environments, for example, in *Pseudomonas putida* when compared to wild-type strains (Compeau et al., 1998).

Continuing the exploration of the culture-independent resistome, the majority of resistance genes identified in this study conferred rifamycin resistance, including *rpoB2, RbpA,* and *HelR*. These genes, along with glycopeptide resistance regulator genes (*vanR/S*), were among the most abundant ARGs in both bulk and plastisphere soils across diverse land uses, as reported in soil resistome surveys from Brazil and China (Ordine et al., 2023; Yang et al., 2022). Although *oleC* was detected in this study, the more commonly reported macrolide efflux homolog is *oleB* (Lawther et al., 2022), suggesting possible variation in efflux gene distribution across environments. Additionally, struvites (Wang et al., 2020) have previously been associated with tetracycline resistance, including containing both tetracycline antibiotic compounds and tetracycline ARGs. However, we did not observe the presence of any tetracycline resistance genes in fertilised soils. Consistent with this, no culturable tetracycline resistance was detected in XIA fertilised soils using the techniques employed here.

Struvites, including those derived from wastewater, have previously been identified as potential reservoirs of ARGs. For example, Chen et al. (2017), using the same XIA piggery struvite as in this study, reported the detection of 165 ARGs and 10 mobile genetic elements (MGEs) in both struvite fertiliser and soil samples. Among these were 30 ARGs deemed high risk and were considered “genes of concern” by the authors, due to their presence in struvite and the phyllosphere but absence in untreated soil. In contrast, we observed a fewer number of ARGs. Although the same XIA piggery struvite source was used, direct comparison remains challenging due to differences in soil pH, methods of ARG identification, different bioinformatic methods and variability in the antibiotic content of the struvite. Additionally, variability in struvite antibiotic content may reflect inconsistencies in production techniques and the heterogeneity of piggery wastewater inputs.

The ARG content of struvites can be influenced by multiple factors, including particle size. Struvite crystals are capable of adsorbing large quantities of extracellular ARGs (eARGs), up to approximately 1.76 × 10¹³ copies per gram, with adsorption efficiency increasing with needle-like crystal shapes and greater surface area (Liao et al., 2025). The primary mechanisms of adsorption include electrostatic interactions and covalent binding, facilitated by magnesium ions and the phosphate backbone of eARGs. Additionally, heavy metals have been shown to promote the migration of antibiotics from wastewater into struvite, although their impact on ARG transport remains inconsistent. Mineral content and organic matter were also found to have no significant effect on ARG mobility (Cai et al., 2020). Another study reported that a majority of ARGs (64.5–94.8%) were associated with high-density suspended solids, which played a key role in their co-precipitation into struvite (Bai et al., 2023).

Superphosphate fertilisers have previously been shown to introduce trace elements into soil, including heavy metals (Heydari et al., 2022). When applied, these elements can accumulate and contribute to elevated levels of AMR, as they co-select for resistance mechanisms such as enhanced efflux and the co-regulation of resistance genes (Heydari et al., 2022). While inorganic fertilisers like TSP have been reported to enrich ARGs in soil, their impact appears to be less pronounced than that of organic fertilisers (Nõlvak et al., 2016). However, in this study, no significant differences in the soil resistome were observed across treatments with TSP, XIA, CG, or untreated soil, as determined by our metagenomic analysis. Similarly, no significant changes were detected in microbial community composition among the treatments.

Finally, we also demonstrated that resistome profiles are highly dependent on the choice of database, read mapping parameters, and cut-off thresholds including gene coverage and the minimum number of reads mapping to a gene to be considered “present”. In contrast, studies employing less stringent or inconsistent criteria than those employed here may report inflated ARG counts. Complementing our findings of low ARG abundance, previous research has shown that when struvite is applied to soil in combination with biochar, the abundance of several ARGs, including *tetX, ermB, sulI*, and *intI1*, can be suppressed by up to 99.95%, with an overall reduction in total ARGs of approximately 30.7% (Li et al., 2020). These results suggest that, when the heavy metal content of struvites is properly managed, they may serve not as a risk factor but instead as a beneficial amendment for reducing both heavy metal toxicity (e.g., copper) and the spread of ARGs in manure-treated soils (Li et al., 2020).

### Struvite application promotes microbial growth without direct selection for Antimicrobial Resistance

Here, we present the first description of how struvite derived from piggery wastewater influences the culturable soil resistome. To the best of our knowledge, this is the first study to explore this impact using culture-based techniques. In contrast to previous research that primarily employed culture-independent methods, predominantly qPCR, our approach pushes beyond this and includes the identification of cultivable isolates (Chen et al., 2017; Li et al., 2019; Li et al., 2020; Wang et al., 2022; Liao et al., 2025).

Our data shows that the XIA struvite treatment generally promoted higher bacterial counts across all conditions and timepoints, suggesting that struvite fertilisation enriches overall microbial growth rather than selecting directly for AMR bacteria. This broad microbial stimulation aligns with trends observed in organic agriculture-derived fertilisers, which are known to support diverse and active microbial communities. For example, El-Mogy et al. (2020) reported that various organic fertilisers significantly increased total nitrogen, organic carbon, and microbial populations, including both fungi and bacteria, compared to mineral fertilisers, with rabbit and chicken manure enhancing both soil fertility and crop yield. Similarly, other studies have also confirmed that organic amendments, such as farmyard manure and pig slurry, promote higher microbial biomass and activity than mineral fertilisation alone (Chakraborty et al., 2011; Ros et al., 2007; Zhong et al., 2010). These effects are likely driven by the high organic carbon content and diverse nutrient profile of organic materials, which fuel microbial growth, improve soil structure, and create a more favourable environment for microbial proliferation (Cui et al., 2023). Therefore, the microbial enrichment observed under the XIA treatment could reflect a similar mechanism to that seen in organic systems, reinforcing the role of nutrient-rich amendments in shaping soil microbial biomass and potentially influencing AMR dynamics.

### Struvite fertilised soil harbours taxonomically and a functionally diverse pool of AMR bacteria

In this study, *Pseudomonas* emerged as the most abundant genus among isolates from XIA treated soil, displaying resistance to multiple antibiotic classes including florfenicol, erythromycin, trimethoprim, and streptomycin. This is a diverse genus of species that occupy a range of ecological niches. For example, *Pseudomonas aeruginosa* is a well-known human pathogen, while species such as *Pseudomonas fluorescens* and *Pseudomonas chlororaphis* are recognised for their roles in promoting plant growth (Ha and Denver, 2018; Tran et al., 2017; Fakhar et al., 2022). Prior studies have reported increased abundance of *Pseudomonas* in urine-derived struvite treatments (Woldeyohannis and Desta, 2024), and streptomycin-resistant strains have been isolated from agricultural soils (Jensen et al., 2001).

This broad resistance aligns with previous findings highlighting the genus’ intrinsic and adaptive resistance mechanisms, particularly in environmental contexts (Silverio et al., 2022; Malik and Aleem, 2011). *Pseudomonas* spp. are motile, non-spore-forming, Gram-negative bacteria commonly found in soil, water, and clinical environments (Hesse et al., 2018; Wisplinghoff, 2017; Fakhar et al., 2022). Their resistance is driven by mechanisms such as efflux pumps, enzymatic inactivation, and membrane modifications (Venter et al., 2015). Notably, *Pseudomonas aeruginosa* strains from soil and compost have been shown to closely resemble clinical isolates, indicating a potential risk of environmental ARG reservoirs contributing to human health threats (Green et al., 1974; Kaszab et al., 2021).

*Bacillus* spp. were the second most abundant genera isolated from struvite-treated soil and, similarly to *Pseudomonas*, showed resistance to 4 antibiotics of different classes, including trimethoprim (5 isolates), amoxicillin (3), erythromycin (2), and neomycin (2), all of which are used as growth promoters, prophylaxis and/or treatment of bacterial infections in livestock farming. Erythromycin resistant *Bacillus cereus* has been previously reported at low levels in agricultural soils (Jensen et al., 2001). *Bacillus* is a resilient, spore-forming, Gram-positive genus with wide environmental distribution, from soil and hot springs to marine sediments and animal guts (Hatamoto et al., 2017; Alina et al., 2015; Soltani et al., 2019). Their robust stress tolerance extends to heavy metals, making them effective agents for biosorption and bioremediation (Wong, 2015; Fakhar et al., 2022). A key environmental stressor influencing ARG prevalence is bioavailable copper, elevated bio-Cu levels were correlated with both class 1 integrons *(intI1)* abundance and ARG prevalence, suggesting that metal stress can drive co-selection for resistance traits (Knapp et al., 2011; Berg et al., 2010; Seiler and Berendonk., 2012). However, Li et al., 2020 also demonstrated that biochar-supported struvite reduced Cu bioavailability and, in turn, ARG selection pressure. In addition to their potential to harbour ARGs, *Bacillus* spp. are widely recognised as plant growth-promoting rhizobacteria (PGPR), enhancing nutrient uptake and crop resistance to pathogens (Shafi et al., 2017). Thus, while they contribute positively to soil health, they may also play a dual role in ARG dynamics, particularly under heavy metal or nutrient-rich fertiliser exposure.

These findings suggest that XIA application supports a taxonomically and functionally diverse reservoir of AMR bacteria, with several genera acting as potential resistance hosts within the soil microbiome. Notably, the isolated genera are commonly associated with natural soil environments and no genera isolated here were obviously associated with only piggery or manure-related sources. However, it remains unclear whether these microbes were specifically selected for by struvite fertilisation, or if they are simply part of the native soil resistome, given that many of them are common soil inhabitants known to exhibit multidrug resistance (MDR).

### Resistome profiles can be driven by tool, database, and parameter choices

When using DeepARG, two key issues became evident in the reported resistome: first, most of the ARGs detected within the soil metagenomes had very low coverage, and second, only a small number of reads were mapped to these ARGs. This low coverage means that, by default, tools like DeepARG can report an ARG as fully present in a sample, even if only short and sometimes non-contiguous fragments are detected. Moreover, many ARGs (and even non-ARG genes) can share highly conserved domains or motifs, meaning that short segments matching an ARG may in fact represent common structural or functional regions found in unrelated genes. As a result, simply detecting partial or fragmented matches is insufficient to confirm the true presence of an ARG in the sample. DeepARG exemplifies a broader issue common in bioinformatics tools: a lack of clarity and transparency in parameter definitions and output metrics. For instance, the precise meaning of reported values like coverage, alignment length, and the alignment start/end positions remains ambiguous, a concern that has been raised on DeepARG’s GitHub (issues page but has yet to be resolved at the time of writing).

Additionally, there is uncertainty surrounding the information provided in DeepARG’s various output files. For example, the .ARG output file (clean.deeparg.mapping.ARG) is described in the GitHub manual (github.com/gaarangoa/deeparg/blob/main/README.md) as including QUERY_END and QUERY_ID. However, this conflicts with the .align.daa file, which is said to follow the BLAST outfmt 6 TAB format. In reality, the query-start and query-end fields in the .ARG file seem to correspond to the sstart and ssend columns of the .align.daa file, indicating that query and subject information have been confused between the two output files. This inconsistency makes interpretation challenging for users. Confusing headers and mismatches between these files have been reported on GitHub but remain unresolved. Additionally, DeepARG does not report gene coverage, therefore we had to calculate this metric ourselves, making it difficult to assess the quality of alignments and potentially leaving users unaware of low-coverage detections, which are described below.

To ensure the detection of ARGs is restricted to nearly complete genes and thereby increasing confidence in their true presence, coverage thresholds are required. This strategy was applied in our alternative read-mapping approach by incorporating both ARG coverage requirements and minimum read thresholds. Furthermore, when using this approach, we observed that the number of ARGs reported was higher under the default Bowtie2 parameters compared with the very sensitive local alignment option. The increased number of reads mapped to ARGs when using the --vsl alignment may reflect the greater sequence diversity for certain ARGs present in soil that are not captured and go unreported under default mapping parameters. This difference could stem from the fact that existing databases are primarily based on clinical isolate data, which may not fully represent the diversity found in environmental resistomes, such as those present in soil samples.

Many of the issues raised here, such as inconsistent ARG coverage reporting, ambiguous output file formats, and unvalidated parameter settings, reflect broader challenges in the development and use of ARG detection tools. These issues are not unique to ARG analysis but apply more widely across bioinformatics, underscoring the need for improved tool development, transparent maintenance, and, crucially, more critical application by users. As Galhano et al., (2021) point out, the lack of standardisation in ARG detection bioinformatics pipelines directly affects the reliability of results. They can also lead to overreporting if the underlying tool outputs are not properly validated, and used without appropriate caution and data exploration, with tools often being used blind. Therefore, conclusions drawn from such analyses may overinflate ARG presence and potentially misdirect future research.

## Conclusions

In conclusion, the impact of struvite fertiliser on the soil resistome and the ARGs associated with struvite fertiliser did not pose a significant AMR risk. Here we propose that struvite fertilisers, such as XIA produced from piggery wastewater, do not selectively facilitate the distribution of ARGs or enhance the growth of AMR bacteria but instead promote bacterial growth. A number of factors could influence the impact struvite has on the soil resistome, including the antibiotic residues with the struvite, the production method, the source of the struvite, as well as the soil and other environmental properties including seasonality. Such variation in these factors could lead to the difference in reports when investigating AMR risk associated with struvites, including those that reported AMR risks associated with struvite fertilisation (Chen *et al.,* 2017; An *et al.,* 2018). Crystal Green and other struvite-based fertilisers could not only act as a potential low AMR risk fertiliser but also address problems such as the P supply chain risk, provide a potential effective solution to P pollution, and offset the mining of new P reserves through recycling thereby creating a sustainable P cycle.

## Data availability

Metagenomic data is available at PRJEB96239 and 16S rRNA gene data is available at PRJEB97999, deposited in the European Nucleotide Archive.

Code used for analysis is available at https://github.com/lawkj/Soil_metagenomic_16S and https://github.com/TheHuwsLab.

## Author Contributions

PNW, RLH-Plant pot experiment design and execution, manuscript review. BG-mass spectrometry analysis, manuscript review. JC-manuscript review. JWM-project supervision, manuscript review. NJD-bioinformatic method development, 16S rRNA gene isolate analysis, manuscript preparation and review. SH-project supervision, manuscript preparation and review. KL-sample collection, methodology, bioinformatic method development, and analysis, laboratory analysis, data curation, manuscript preparation, and review.

## Acknowledgements.

Authors would like to thank Dr Marina Reyne and the Queen’s University Belfast Wastewater Epidemiology team for their technical assistance with the Nanopore GridIONxs sequencing. The authors would also like to thank Dr William O’Neill for his help with DNA extracting from the struvite fertiliser.

## Supplementary material

**Supplementary Figure 1:**
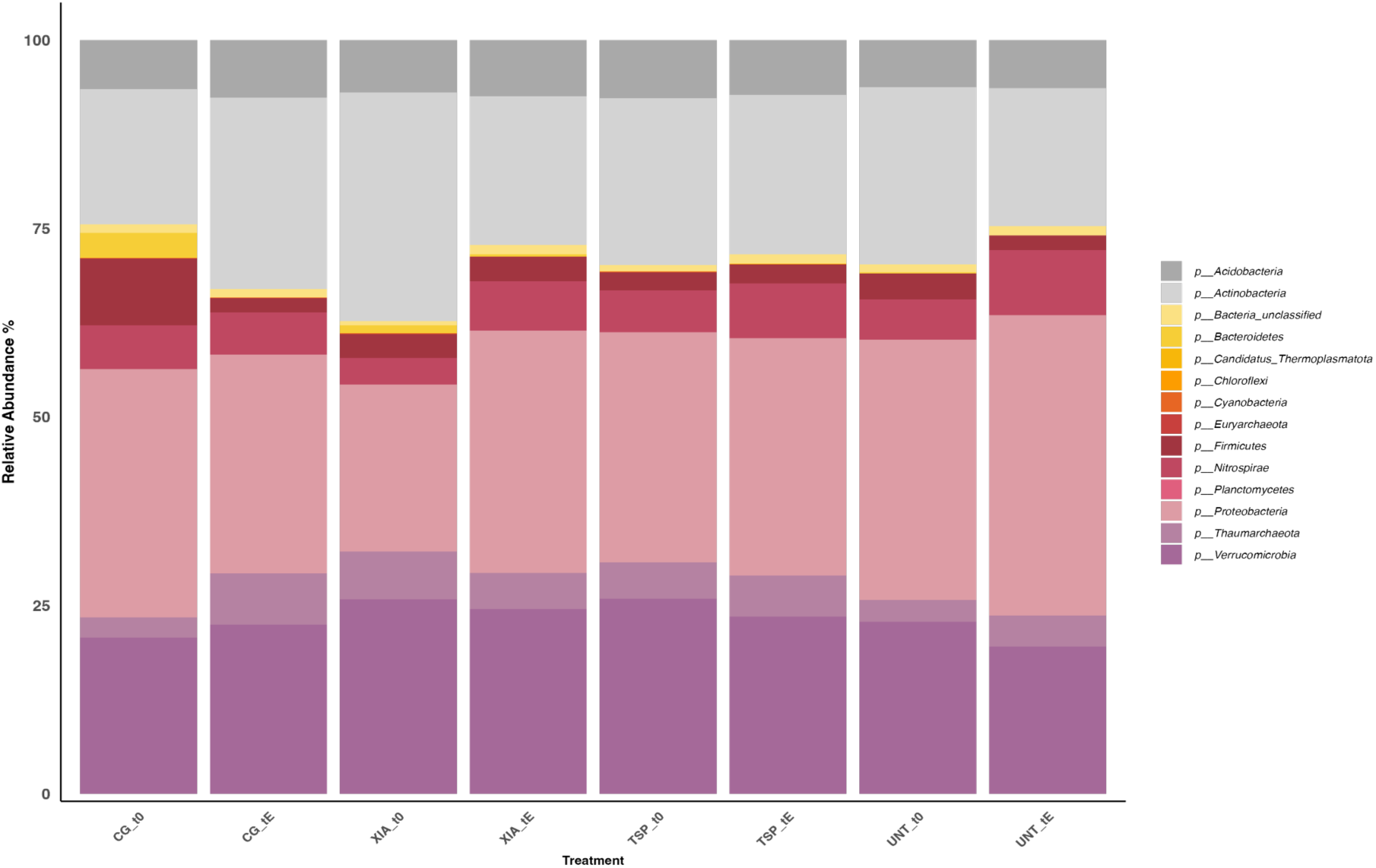
Taxonomic composition of soil metagenomic reads assigned using MetaPhlAn. Stacked bar plots display the relative abundance of the top 20 most abundant phyla across different treatments and two time points (t0 = time zero, tE = end time point). Phyla outside the top 20 are grouped as "Other" for clarity. Treatments include: UNT (Untreated), CG (Crystal Green), XIA (Xiamen), and TSP (Triple Super Phosphate).

**Supplementary Table 1:** Full LC-MS/MS results from soil and fertiliser samples.

**Supplementary Table 2:** List of antibiotics detectable by LC-MS/MS (Percentage recovered (%)).

**Supplementary Table 3:** MetaPhlAn genus level classification control samples: negative “kitome” control and positive (Zymo mock community) compared to expected composition.

**Supplementary Table 4:** MetaPhlAn results for soil metagenome samples at the genus level.

**Supplementary Table 5:** Taxonomic diversity statistics across soil metagenomes (genus, phylum, Simpson, Shannon, Jaccard and Bray Curtis analysis).

**Supplementary Table 6:** DeepARG complete results for soil metagenomics including default parameters and a --gene_coverage cutoff of 0.2.

**Supplementary Table 7:** ARGs identified in soil metagenomes using default DeepARG parameters and computed (inhouse) gene converge.

**Supplementary Table 8:** ARGs identified using a readmapping approach (Bowtie2) using very_sensitive_local and all alignments reported, then a coverage cut of 90% and a minimum number of reads of 100 applied.

**Supplementary Table 9:** EMU genus level classification of two positive control samples (Zymo mock community) compared to expected composition.

## References

1. Ågren, G.I., Wetterstedt, J.M. and Billberger, M.F., 2012. Nutrient limitation on terrestrial plant growth–modeling the interaction between nitrogen and phosphorus. New Phytologist, 194(4), pp.953–960.

2. Alcock, B.P., Huynh, W., Chalil, R., Smith, K.W., Raphenya, A.R., Wlodarski, M.A., Edalatmand, A., Petkau, A., Syed, S.A., Tsang, K.K. and Baker, S.J., 2023. CARD 2023: expanded curation, support for machine learning, and resistome prediction at the Comprehensive Antibiotic Resistance Database. Nucleic acids research, 51(D1), pp.D690–D699.

3. Alina, S.O., Constantinscu, F. and Petruţa, C.C., 2015. Biodiversity of Bacillus subtilis group and beneficial traits of Bacillus species useful in plant protection. Romanian Biotechnological Letters, 20(5), pp.10737–10750.

4. An, X.L., Chen, Q.L., Zhu, D. and Su, J.Q., 2018. Distinct effects of struvite and biochar amendment on the class 1 integron antibiotic resistance gene cassettes in phyllosphere and rhizosphere. Science of The Total Environment, 631, pp.668–676.

5. Ar, Y., 2009. The disappearing nutrient. Nature, 461, p.8.

6. Arango-Argoty, G., Garner, E., Pruden, A., Heath, L.S., Vikesland, P. and Zhang, L., 2018. DeepARG: a deep learning approach for predicting antibiotic resistance genes from metagenomic data. Microbiome, 6(1), p.23.

7. Bai, W., Tang, R., Wu, G., Wang, W., Yuan, S., Xiao, L., Zhan, X. and Hu, Z.H., 2023. Role of suspended solids on the co-precipitation of pathogenic indicators and antibiotic resistance genes with struvite from digested swine wastewater. Journal of Hazardous Materials, 459, p.132235.

8. Berg, J., Thorsen, M.K., Holm, P.E., Jensen, J., Nybroe, O. and Brandt, K.K., 2010. Cu exposure under field conditions coselects for antibiotic resistance as determined by a novel cultivation-independent bacterial community tolerance assay. Environmental science & technology, 44(22), pp.8724–8728.

9. Blanco-Míguez, A., Beghini, F., Cumbo, F., McIver, L.J., Thompson, K.N., Zolfo, M., Manghi, P., Dubois, L., Huang, K.D., Thomas, A.M. and Nickols, W.A., 2023. Extending and improving metagenomic taxonomic profiling with uncharacterized species using MetaPhlAn 4. Nature Biotechnology, 41(11), pp.1633–1644.

10. Blank, L.M., 2012. The cell and P: from cellular function to biotechnological application. Current opinion in biotechnology, 23(6), pp.846–851.

11. Bolger, A.M., Lohse, M. and Usadel, B., 2014. Trimmomatic: a flexible trimmer for Illumina sequence data. Bioinformatics, 30(15), pp.2114–2120.

12. Bortolaia, V., Kaas, R.S., Ruppe, E., Roberts, M.C., Schwarz, S., Cattoir, V., Philippon, A., Allesoe, R.L., Rebelo, A.R., Florensa, A.F. and Fagelhauer, L., 2020. ResFinder 4.0 for predictions of phenotypes from genotypes. Journal of Antimicrobial Chemotherapy, 75(12), pp.3491–3500.

13. Buchfink, B., Reuter, K. and Drost, H.G., 2021. Sensitive protein alignments at tree-of-life scale using DIAMOND. Nature methods, 18(4), pp.366–368.

14. Cai, J., Ye, Z.L., Ye, C., Ye, X. and Chen, S., 2020. Struvite crystallization induced the discrepant transports of antibiotics and antibiotic resistance genes in phosphorus recovery from swine wastewater. Environmental Pollution, 266, p.115361.

15. Chakraborty, A., Chakrabarti, K., Chakraborty, A. and Ghosh, S., 2011. Effect of long-term fertilizers and manure application on microbial biomass and microbial activity of a tropical agricultural soil. Biology and fertility of Soils, 47, pp.227–233.

16. Chen, Q.L., An, X.L., Zhu, Y.G., Su, J.Q., Gillings, M.R., Ye, Z.L. and Cui, L., 2017. Application of struvite alters the antibiotic resistome in soil, rhizosphere, and phyllosphere. Environmental Science & Technology, 51(14), pp.8149–8157.

17. Compeau, G., Al-Achi, B.J., Platsouka, E. and Levy, S.B., 1988. Survival of rifampin-resistant mutants of Pseudomonas fluorescens and Pseudomonas putida in soil systems. Applied and Environmental Microbiology, 54(10), pp.2432–2438.

18. Cordell, D., 2008. The Story of Phosphorus: missing global governance of a critical resource. SENSE Earth Systems Governance, Amsterdam.

19. Cordell, D. and White, S., 2011. Peak phosphorus: clarifying the key issues of a vigorous debate about long-term phosphorus security. Sustainability, 3(10), pp.2027–2049.

20. Cordell, D. and White, S., 2013. Sustainable phosphorus measures: strategies and technologies for achieving phosphorus security. Agronomy, 3(1), pp.86–116.

21. Cui, J., Yang, B., Zhang, M., Song, D., Xu, X., Ai, C., Liang, G. and Zhou, W., 2023. Investigating the effects of organic amendments on soil microbial composition and its linkage to soil organic carbon: A global meta-analysis. Science of The Total Environment, 894, p.164899.

22. Curry, K.D., Wang, Q., Nute, M.G., Tyshaieva, A., Reeves, E., Soriano, S., Wu, Q., Graeber, E., Finzer, P., Mendling, W. and Savidge, T., 2022. Emu: species-level microbial community profiling of full-length 16S rRNA Oxford Nanopore sequencing data. Nature methods, 19(7), pp.845–853.

23. Danecek, P., Bonfield, J.K., Liddle, J., Marshall, J., Ohan, V., Pollard, M.O., Whitwham, A., Keane, T., McCarthy, S.A., Davies, R.M. and Li, H., 2021. Twelve years of SAMtools and BCFtools. Gigascience, 10(2), p.giab008.

24. De Boer, M.A., Kabbe, C. and Slootweg, J.C., 2018. Comment on “Application of Struvite Alters the Antibiotic Resistome in Soil, Rhizosphere, and Phyllosphere”. Environmental science & technology, 52(24), pp.14564–14565.

25. De Coster, W. and Rademakers, R., 2023. NanoPack2: population-scale evaluation of long-read sequencing data. Bioinformatics, 39(5), p.btad311.

26. Degryse, F., Baird, R., Da Silva, R.C. and McLaughlin, M.J., 2017. Dissolution rate and agronomic effectiveness of struvite fertilisers–effect of soil pH, granulation and base excess. Plant and soil, 410(1-2), pp.139–152.

27. Dixon, P., 2003. VEGAN, a package of R functions for community ecology. Journal of vegetation science, 14(6), pp.927–930.

28. Doran, J.L., Pang, Y., Mdluli, K.E., Moran, A.J., Victor, T.C., Stokes, R.W., Mahenthiralingam, E., Kreiswirth, B.N., Butt, J.L., Baron, G.S. and Treit, J.D., 1997. Mycobacterium tuberculosis efpA encodes an efflux protein of the QacA transporter family. Clinical Diagnostic Laboratory Immunology, 4(1), pp.23–32

29. El-Mogy, M.M., Abdelaziz, S.M., Mahmoud, A.W.M., Elsayed, T.R., Abdel-Kader, N.H. and Mohamed, M.I., 2020. Comparative effects of different organic and inorganic fertilisers on soil fertility, plant growth, soil microbial community, and storage ability of lettuce. Agriculture, 66(3), pp.87–107

30. Etter, B., Tilley, E., Khadka, R. and Udert, K.M., 2011. Low-cost struvite production using source-separated urine in Nepal. Water research, 45(2), pp.852–862.

31. European Commission, 2018. Antibiotic Resistance in Struvite Fertiliser from Waste Water Could Enter the Food Chain. Sci. for Environ. Policy.

32. European Commission, 2021. Fertilising products – precipitated phosphate salts and derivates. [online] Available at: https://ec.europa.eu/info/law/better-regulation/have-your-say/initiatives/12163-Fertilising-products-precipitated-phosphate-salts-and-derivates_en [Accessed 07 Jan 2024].

33. European Commission. 2020. Communication from the Commission to the European Parliament, the Council, the European Economic and Social Committee and the Committee of the regions. Critical Raw Materials Resilience: Charting a Path towards greater Security and Sustainability. COM/2020/4.

34. European Medicines Agency (EMA), 2019. Quinolone- and fluoroquinolone-containing medicinal products. [online] European Medicines Agency. Available at: https://www.ema.europa.eu/en/medicines/human/referrals/quinolone-fluoroquinolone-containing-medicinal-products [Accessed 1 July 2025].

35. Everaert, M., da Silva, R.C., Degryse, F., McLaughlin, M.J. and Smolders, E., 2018. Limited dissolved phosphorus runoff losses from layered double hydroxide and struvite fertilisers in a rainfall simulation study. Journal of environmental quality, 47(2), pp.371–377.

36. Evoqua Water Technologies, 2021. OSTARA’S PEARL® SYSTEM BY EVOQUA: The standard in nutrient recovery [Online]. Available at: <https://www.evoqua.com/siteassets/documents/products/anaerobic/mu-ostarapearl-br-0222-lores.pdf> [accessed 6 December 2022].

37. Fakhar, A., Gul, B., Gurmani, A.R., Khan, S.M., Ali, S., Sultan, T., Chaudhary, H.J., Rafique, M. and Rizwan, M., 2022. Heavy metal remediation and resistance mechanism of Aeromonas, Bacillus, and Pseudomonas: A review. Critical Reviews in Environmental Science and Technology, 52(11), pp.1868–1914.

38. Felsenstein, J., 1993. Distributed by the author. Phylogeny Inference Package, Version 3.5 c.

39. Galhano, B.S., Ferrari, R.G., Panzenhagen, P., de Jesus, A.C.S. and Conte-Junior, C.A., 2021. Antimicrobial resistance gene detection methods for bacteria in animal-based foods: A brief review of highlights and advantages. Microorganisms, 9(5), p.923.

40. Gao, D., Li, B., Huang, X., Liu, X., Li, R., Ye, Z., Wu, X., Huang, Y. and Wang, G., 2023. A review of the migration mechanism of antibiotics during struvite recovery from wastewater. Chemical Engineering Journal, 466, p.142983.

41. Green, S.K., Schroth, M.N., Cho, J.J., Kominos, S.D. and Vitanza-Jack, V.B., 1974. Agricultural plants and soil as a reservoir for Pseudomonas aeruginosa. Applied microbiology, 28(6), pp.987–991.

42. Guglierame, P., Pasca, M.R., De Rossi, E., Buroni, S., Arrigo, P., Manina, G. and Riccardi, G., 2006. Efflux pump genes of the resistance-nodulation-division family in Burkholderia cenocepacia genome. BMC microbiology, 6, pp.1–14.

43. Ha, A.D. and Denver, D.R., 2018. Comparative genomic analysis of 130 bacteriophages infecting bacteria in the genus Pseudomonas. Frontiers in microbiology, 9, p.1456.

44. Hall, R.L., Staal, L.B., Macintosh, K.A., McGrath, J.W., Bailey, J., Black, L., Nielsen, U.G., Reitzel, K. and Williams, P.N., 2020. Phosphorus speciation and fertiliser performance characteristics: A comparison of waste recovered struvites from global sources. Geoderma, 362, p.114096.

45. Hashemzadeh, M., Hasanvand, M. and Montazeri, E.A., 2025. Analysis of relative genes expression and mutation of pstB and efpA efflux pumps in Mycobacterium simiae isolates from suspected tuberculosis patients by using quantitative Real-time PCR. BMC microbiology, 25(1), p.144.

46. Hatamoto, M., Kaneko, T., Takimoto, Y., Ito, T., Miyazato, N., Maki, S., Yamaguchi, T. and Aoi, T., 2017. Microbial community structure and enumeration of Bacillus species in activated sludge. Journal of Water and Environment Technology, 15(6), pp.233–240.

47. Hesse, C., Schulz, F., Bull, C.T., Shaffer, B.T., Yan, Q., Shapiro, N., Hassan, K.A., Varghese, N., Elbourne, L.D., Paulsen, I.T. and Kyrpides, N., 2018. Genome-based evolutionary history of Pseudomonas spp. Environmental microbiology, 20(6), pp.2142–2159.

48. Heuer, H. and Smalla, K., 2007. Manure and sulfadiazine synergistically increased bacterial antibiotic resistance in soil over at least two months. Environmental Microbiology, 9(3), pp.657–666.

49. Heuer, H., Schmitt, H. and Smalla, K., 2011. Antibiotic resistance gene spread due to manure application on agricultural fields. Current opinion in microbiology, 14(3), pp.236–243.

50. Heydari, A., Kim, N.D., Horswell, J., Gielen, G., Siggins, A., Taylor, M., Bromhead, C. and Palmer, B.R., 2022. Co-Selection of Heavy Metal and Antibiotic Resistance in Soil Bacteria from Agricultural Soils in New Zealand. Sustainability, 14(3), p.1790.

51. Jasinski, S.M., 2021. Mineral commodity summaries: phosphate rock. US Geological Survey.

52. Jensen, L.B., Baloda, S., Boye, M. and Aarestrup, F.M., 2001. Antimicrobial resistance among Pseudomonas spp. and the Bacillus cereus group isolated from Danish agricultural soil. Environment international, 26(7-8), pp.581–587.

53. Kaszab, E., Radó, J., Kriszt, B., Pászti, J., Lesinszki, V., Szabó, A., Tóth, G., Khaledi, A. and Szoboszlay, S., 2021. Groundwater, soil and compost, as possible sources of virulent and antibiotic-resistant Pseudomonas aeruginosa. International Journal of Environmental Health Research, 31(7), pp.848–860.

54. Kenjeric, L., Sulyok, M., Malachova, A., Greer, B., Kolawole, O., Quinn, B., Elliott, C.T. and Krska, R., 2024. Extention and interlaboratory comparison of an LC-MS/MS multi-class method for the determination of 15 different classes of veterinary drug residues in milk and poultry feed. Food Chemistry, 449, p.138834.

55. Khurana, P., Pulicharla, R. and Brar, S.K., 2021. Antibiotic-metal complexes in wastewaters: fate and treatment trajectory. Environment International, 157, p.106863.

56. Kisil, O.V., Efimenko, T.A. and Efremenkova, O.V., 2021. Looking back to Amycolatopsis: history of the antibiotic discovery and future prospects. Antibiotics, 10(10), p.1254.

57. Kleinman, P., Sharpley, A., Buda, A., McDowell, R. and Allen, A., 2011. Soil controls of phosphorus in runoff: Management barriers and opportunities. Canadian Journal of Soil Science, 91(3), pp.329–338.

58. Knapp, C.W., McCluskey, S.M., Singh, B.K., Campbell, C.D., Hudson, G. and Graham, D.W., 2011. Antibiotic resistance gene abundances correlate with metal and geochemical conditions in archived Scottish soils. PloS one, 6(11), p.e27300.

59. Langmead, B. and Salzberg, S.L., 2012. Fast gapped-read alignment with Bowtie 2. Nature methods, 9(4), pp.357–359.

60. Lawther, K., Santos, F.G., Oyama, L.B., Rubino, F., Morrison, S., Creevey, C.J., McGrath, J.W. and Huws, S.A., 2022. Resistome analysis of global livestock and soil microbiomes. Frontiers in microbiology, 13, p.897905.

61. Li, Y., Wang, X., Li, J., Wang, Y., Song, J., Xia, S., Jing, H. and Zhao, J., 2019. Effects of struvite-humic acid loaded biochar/bentonite composite amendment on Zn (II) and antibiotic resistance genes in manure-soil. Chemical Engineering Journal, 375, p.122013.

62. Li, Y., Wang, X., Wang, Y., Wang, F., Xia, S. and Zhao, J., 2020. Struvite-supported biochar composite effectively lowers Cu bio-availability and the abundance of antibiotic-resistance genes in soil. Science of the Total Environment, 724, p.138294.

63. Liao, W., Huang, X., Ye, Z.L., Zhang, T., Cai, J., Huang, Y. and Li, Y., 2025. Insights into the migration mechanism of extracellular antibiotic resistance genes during struvite recovery using synthetic wastewater. Water Research, 268, p.122681.

64. Lou, Y., Ye, X., Ye, Z.L., Chiang, P.C. and Chen, S., 2018. Occurrence and ecological risks of veterinary antibiotics in struvite recovered from swine wastewater. Journal of cleaner production, 201, pp.678–685.

65. Malik, A. and Aleem, A., 2011. Incidence of metal and antibiotic resistance in Pseudomonas spp. from the river water, agricultural soil irrigated with wastewater and groundwater. Environmental Monitoring and Assessment, 178, pp.293–308.

66. Mar, S.S. and Okazaki, M., 2012. Investigation of Cd contents in several phosphate rocks used for the production of fertilizer. Microchemical Journal, 104, pp.17–21.

67. Martinez Arbizu, P. (2020). pairwiseAdonis: Pairwise multilevel comparison using adonis. R package version 0.4

68. Mima, T., Kohira, N., Li, Y., Sekiya, H., Ogawa, W., Kuroda, T. and Tsuchiya, T., 2009. Gene cloning and characteristics of the RND-type multidrug efflux pump MuxABC-OpmB possessing two RND components in Pseudomonas aeruginosa. Microbiology, 155(11), pp.3509–3517.

69. Muys, M., Phukan, R., Brader, G., Samad, A., Moretti, M., Haiden, B., Pluchon, S., Roest, K., Vlaeminck, S.E. and Spiller, M., 2021. A systematic comparison of commercially produced struvite: Quantities, qualities and soil-maize phosphorus availability. Science of the total Environment, 756, p.143726.

70. Niño-Savala, A.G., Zhuang, Z., Ma, X., Fangmeier, A., Li, H., Tang, A. and Liu, X., 2019. Cadmium pollution from phosphate fertilizers in arable soils and crops: an overview. Front. Agric. Sci. Eng, 6, pp.419–430.

71. Nõlvak, H., Truu, M., Kanger, K., Tampere, M., Espenberg, M., Loit, E., Raave, H. and Truu, J., 2016. Inorganic and organic fertilizers impact the abundance and proportion of antibiotic resistance and integron-integrase genes in agricultural grassland soil. Science of the Total Environment, 562, pp.678–689.

72. Oelkers, E.H. and Valsami-Jones, E., 2008. Phosphate mineral reactivity and global sustainability. Elements, 4(2), pp.83–87.

73. Ohtake, H. and Tsuneda, S. eds., 2019. Phosphorus recovery and recycling (p. 2019). Singapore: Springer Singapore.

74. Ordine, J.V.W., De Souza, G.M., Tamasco, G., Virgilio, S., Fernandes, A.F.T., Silva-Rocha, R. and Guazzaroni, M.E., 2023. Metagenomic insights for antimicrobial resistance surveillance in soils with different land uses in Brazil. Antibiotics, 12(2), p.334.

75. Ostara, 2021, From struvite to stewardship [Online]. Available at: https://ostara.com/municipal-utilities/ [Accessed 5 April 2022].

76. Ott, C. and Rechberger, H., 2012. The European phosphorus balance. Resources, conservation and recycling, 60, pp.159–172.

77. Rahman, M.M., Liu, Y., Kwag, J.H. and Ra, C., 2011. Recovery of struvite from animal wastewater and its nutrient leaching loss in soil. Journal of hazardous materials, 186(2-3), pp.2026–2030.

78. Rahman, M.M., Salleh, M.A.M., Rashid, U., Ahsan, A., Hossain, M.M. and Ra, C.S., 2014. Production of slow release crystal fertilizer from wastewaters through struvite crystallization–A review. Arabian journal of chemistry, 7(1), pp.139–155.

79. Reid, N., Reyne, M.I., O’Neill, W., Greer, B., He, Q., Burdekin, O., McGrath, J.W. and Elliott, C.T., 2024. Unprecedented Harmful algal bloom in the UK and Ireland’s largest lake associated with gastrointestinal bacteria, microcystins and anabaenopeptins presenting an environmental and public health risk. Environment international, 190, p.108934.

80. Ros, M., García, C. and Hernandez, M.T., 2007. Evaluation of different pig slurry composts as fertilizer of horticultural crops: Effects on selected chemical and microbial properties. Renewable Agriculture and Food Systems, 22(4), pp.307–315.

81. Schoch, C.L., Ciufo, S., Domrachev, M., Hotton, C.L., Kannan, S., Khovanskaya, R., Leipe, D., Mcveigh, R., O’Neill, K., Robbertse, B. and Sharma, S., 2020. NCBI Taxonomy: a comprehensive update on curation, resources and tools. Database, 2020, p.baaa062.

82. Seiler, C. and Berendonk, T.U., 2012. Heavy metal driven co-selection of antibiotic resistance in soil and water bodies impacted by agriculture and aquaculture. Frontiers in microbiology, 3, p.399.

83. Shafi, J., Tian, H. and Ji, M., 2017. Bacillus species as versatile weapons for plant pathogens: a review. Biotechnology & Biotechnological Equipment, 31(3), pp.446–459

84. Silverio, M.P., Kraychete, G.B., Rosado, A.S. and Bonelli, R.R., 2022. Pseudomonas fluorescens complex and its intrinsic, adaptive, and acquired antimicrobial resistance mechanisms in pristine and human-impacted sites. Antibiotics, 11(8), p.985.

85. Soltani, M., Ghosh, K., Hoseinifar, S.H., Kumar, V., Lymbery, A.J., Roy, S. and Ringø, E., 2019. Genus Bacillus, promising probiotics in aquaculture: aquatic animal origin, bio-active components, bioremediation and efficacy in fish and shellfish. Reviews in Fisheries Science & Aquaculture, 27(3), pp.331–379.

86. Somprasong, N., Yi, J., Hall, C.M., Webb, J.R., Sahl, J.W., Wagner, D.M., Keim, P., Currie, B.J. and Schweizer, H.P., 2021. Conservation of resistance-nodulation-cell division efflux pump-mediated antibiotic resistance in Burkholderia cepacia complex and Burkholderia pseudomallei complex species. Antimicrobial Agents and Chemotherapy, 65(9), pp.10–1128.

87. Tran, P.N., Savka, M.A. and Gan, H.M., 2017. In-silico taxonomic classification of 373 genomes reveals species misidentification and new genospecies within the genus Pseudomonas. Frontiers in Microbiology, 8, p.1296.

88. Udikovic-Kolic, N., Wichmann, F., Broderick, N.A. and Handelsman, J., 2014. Bloom of resident antibiotic-resistant bacteria in soil following manure fertilization. Proceedings of the National Academy of Sciences, 111(42), pp.15202–15207.

89. Uludag-Demirer, S., Demirer, G.N. and Chen, S.J.P.B., 2005. Ammonia removal from anaerobically digested dairy manure by struvite precipitation. Process Biochemistry, 40(12), pp.3667–3674.

90. Venter, H., Mowla, R., Ohene-Agyei, T. and Ma, S., 2015. RND-type drug efflux pumps from Gram-negative bacteria: molecular mechanism and inhibition. Frontiers in microbiology, 6, p.377.

91. Wang, Y., Wang, X., Li, J., Li, Y., Liu, Y., Wang, F. and Zhao, J., 2020. Adsorption and precipitation behaviors of zinc, copper and tetracycline with struvite products obtained by phosphorus recovery from swine wastewater. Journal of Environmental Chemical Engineering, 8(6), p.104488.

92. Wichmann, F., Udikovic-Kolic, N., Andrew, S. and Handelsman, J., 2014. Diverse antibiotic resistance genes in dairy cow manure. MBio, 5(2), pp.e01017–13.

93. Wick R. (2017) Porechop: adapter trimmer for Oxford Nanopore reads. https://github.com/rrwick/Porechop/.

94. Wisplinghoff, H., 2017. Pseudomonas spp., Acinetobacter spp. and miscellaneous Gram-negative bacilli. In Infectious diseases (pp. 1579-1599). Elsevier.

95. Woldeyohannis, N.N. and Desta, A.F., 2023. Fate of antimicrobial resistance genes (ARG) and ARG carriers in struvite production process from human urine. *Journal of Environmental Science and Health*, Part A, 58(9), pp.783–792.

96. Woldeyohannis, N.N. and Desta, A.F., 2024. Metagenome-based microbial community analysis of urine-derived fertilizer. BMC microbiology, 24(1), p.418.

97. Wong, L.S., 2015. Microbial cementation of ureolytic bacteria from the genus Bacillus: a review of the bacterial application on cement-based materials for cleaner production. Journal of Cleaner Production, 93, pp.5–17.

98. Yang, Y., Li, T., Liu, P., Li, H. and Hu, F., 2022. The formation of specific bacterial communities contributes to the enrichment of antibiotic resistance genes in the soil plastisphere. Journal of Hazardous Materials, 436, p.129247

99. Ye, Z.L., Chen, S.H., Wang, S.M., Lin, L.F., Yan, Y.J., Zhang, Z.J. and Chen, J.S., 2010. Phosphorus recovery from synthetic swine wastewater by chemical precipitation using response surface methodology. Journal of Hazardous Materials, 176(1-3), pp.1083–1088.

100. Yu, G., Smith, D.K., Zhu, H., Guan, Y. and Lam, T.T.Y., 2017. ggtree: an R package for visualization and annotation of phylogenetic trees with their covariates and other associated data. Methods in ecology and evolution, 8(1), pp.28–36.

101. Zhong, W., Gu, T., Wang, W., Zhang, B., Lin, X., Huang, Q. and Shen, W., 2010. The effects of mineral fertilizer and organic manure on soil microbial community and diversity. Plant and soil, 326, pp.511–522.

